# Targeted disruption of Oncogenic Biosphere: A Paradigm Shift Beyond Cancer Cell-Centric Therapies

**DOI:** 10.1101/2025.07.20.665731

**Authors:** N Sanoj Rejinold, Sujong Kim, Geun-woo Jin, Jin-Ho Choy

**Author notes:** These authors contributed equally to this work (N Sanoj Rejinold, Sujong Kim).

## Abstract

Traditional cancer therapies have primarily focused on directly targeting malignant cells, yet this paradigm has consistently fallen short in addressing therapeutic resistance and metastasis^1^. These persistent challenges suggest that the tumor’s surrounding environment—not just the cancer cells themselves—plays a decisive role in treatment failure. In this study, we present an ecosystem-oriented strategy that targets the pathological microenvironment sustaining cancer progression. We repurposed niclosamide, an FDA-approved anthelmintic agent^2,3^, and enhanced its bioavailability by formulating Penetrium an orally deliverable nanotherapeutic composed of niclosamide, magnesium oxide, and hydroxypropyl methylcellulose. In syngeneic and xenograft models of triple-negative breast cancer (TNBC) and non-small cell lung cancer (NSCLC), Penetrium showed robust antitumor synergy when combined with standard therapies including anti-PD-1, paclitaxel, and anti-VEGF agents. Notably, Penetrium markedly blocked metastatic spread. Mechanistic analyses revealed that Penetrium remodels the tumor extracellular matrix (ECM), downregulates MMP-9, restores E-cadherin, and enhances immune infiltration— thereby converting immune-excluded tumors into immune-permissive niches. Furthermore, in patient-derived pancreatic tumor organoids co-cultured with cancer-associated fibroblasts (CAFs), we demonstrated that tumor cell survival is critically dependent on the stromal context. Importantly, Penetrium selectively eliminated CAFs without affecting normal fibroblasts. These findings validate the oncogenic biosphere, and especially the CAF-rich stroma, as a critical therapeutic target. Our results underscore a shift from the conventional cancer cell-centric model to an oncogenic biosphere-disruption strategy that dismantles the structural and immunological defenses of desmoplastic cancers, offering a promising avenue for overcoming resistance and metastasis.

---

Despite decades of progress in cancer therapeutics, treatment resistance and metastasis remain formidable challenges across multiple tumor types^1^. These clinical obstacles are particularly pronounced in desmoplastic cancers such as triple-negative breast cancer (TNBC)^4^, pancreatic ductal adenocarcinoma (PDAC)^5^, and metastatic prostate and lung cancers characterized not only by their intrinsic genetic aggressiveness but also by their highly remodeled tumor microenvironments (TMEs)^6-8^. It is increasingly evident that effective cancer therapy must move beyond the traditional cancer cell–centric paradigm and instead target the complex, dynamic oncogenic biosphere that supports malignant progression^9^.

Among the key components of this ecosystem, the extracellular matrix (ECM) has emerged as a critical structural and signaling barrier that regulates therapeutic response, immune cell infiltration, and tumor dissemination^10,11^. In TNBC, which affects approximately 15–20% of breast cancer patients and is associated with rapid progression and early recurrence, the ECM is profoundly dysregulated^12^. The desmoplastic stroma in such tumors is typified by extensive collagen deposition, enhanced matrix stiffness, and abnormal expression of ECM-associated enzymes and glycoproteins^13-22^. These pathological features do not merely serve as passive scaffolds but actively contribute to tumor progression by promoting epithelial–mesenchymal transition (EMT)^23^, facilitating immune exclusion, and reducing intratumoral drug penetration. As a result, even potent cytotoxic and immunotherapeutic agents are rendered ineffective within ECM-rich niches, giving rise to what is now termed “pseudo-resistance”— a resistance mechanism not due to intrinsic drug insensitivity but rather to a hostile tumor architecture^24^.

**Scheme 1.**
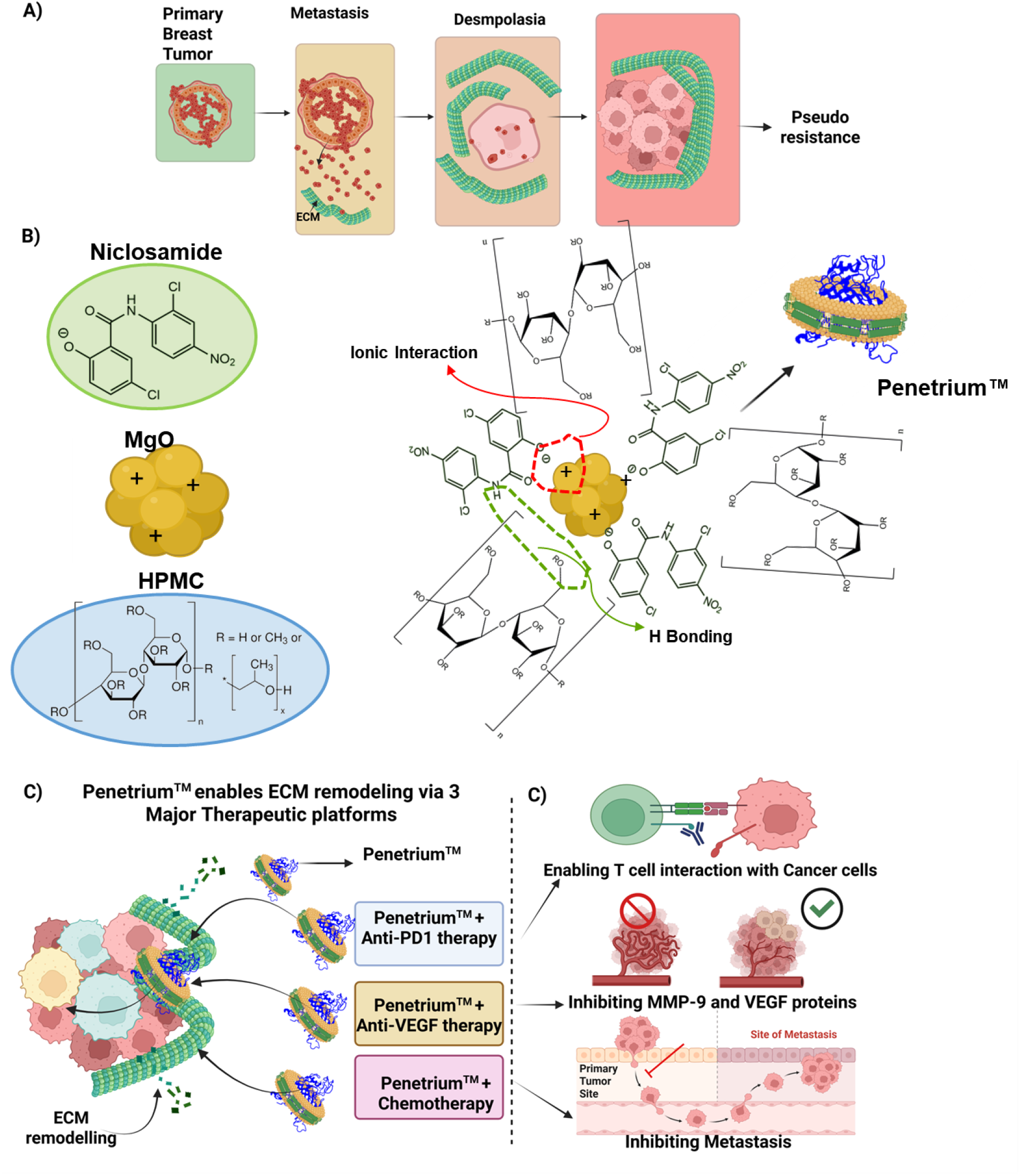
Penetrium Nanodrug Remodels the Tumor ECM to Overcome Pseudo-Resistance and Enhance Anticancer Therapeutic Synergy (A) Schematic representation of the metastatic cascade and the emergence of ECM-mediated pseudo-resistance in triple-negative breast cancer (TNBC). Tumor progression begins with a primary breast tumor that invades the surrounding extracellular matrix (ECM) during metastasis. Disseminated cancer cells colonize distant organs, where they induce desmoplasia, characterized by excessive ECM deposition, including collagen and fibronectin. This fibrotic microenvironment impedes drug diffusion, creating physical and biochemical barriers that lead to pseudo-resistance—a non-mutational treatment failure state; (B) Chemistry of Penetrium where ionic interaction between Niclosamide and MgO are predominant, whereas HOMC is coated by H bonding; C) Mechanism of Penetrium-mediated ECM remodeling and its integration with four major therapeutic platforms. Penetrium is a nanoengineered repurposed niclosamide (NIC) which is bonded on Magnesium oxide (MgO) (via ionic interaction), and hydroxypropyl methylcellulose (HPMC) (via physical coating), originally repurposed from antiviral applications. Designed with ECM-interactive ligands and optimized physicochemical features, Penetrium selectively binds to and disrupts fibrotic ECM components. This remodeling improves drug access and stromal penetration, enabling co-delivery with: Checkpoint inhibitors (Anti-PD1 therapy) to enhance immune recognition, Anti-angiogenic agents (Anti-VEGF therapy) to normalize tumor vasculature, Cytotoxic agents (Chemotherapy) to increase efficacy against previously inaccessible tumor cell populations; (C) Therapeutic outcomes of Penetrium-based stromal modulation. By degrading ECM barriers and inhibiting matrix metalloproteinase-9 (MMP-9) and vascular endothelial growth factor (VEGF) expression, Penetrium facilitates: Enhanced T-cell interaction with tumor cells, improving immunotherapy responses, Suppression of aberrant angiogenesis, leading to normalized vasculature and reduced tumor hypoxia, Inhibition of metastatic dissemination, disrupting the migration of tumor cells from the primary site to distant organs. Overall, Penetrium represents a versatile ECM-modulating nanoplatform capable of reversing stromal-mediated pseudo-resistance and boosting the performance of multiple anticancer therapies.

Cancer-associated fibroblasts (CAFs), a dominant stromal cell population within the TME, are key orchestrators of ECM remodeling^25^. By secreting matrix proteins (e.g., collagen I, fibronectin), crosslinking enzymes (e.g., lysyl oxidase), and immunosuppressive factors, CAFs create a dense and immunologically cold environment that protects tumor cells from both immune attack and therapeutic agents^25^. While conventional approaches have aimed to kill cancer cells directly, accumulating evidence suggests that such strategies often leave the supportive stromal architecture intact, allowing the tumor to re-emerge or disseminate through pre-established survival routes.

Hence, targeting the oncogenic biosphere—particularly its ECM-rich, CAF-dominated stroma—has gained traction as a promising therapeutic avenue. effects extend beyond cancer cells to target the structural and immunological barriers that define the desmoplastic tumor niche.

In this study, we investigate the oncogenic biosphere-modulating effects of Penetrium in preclinical models of TNBC and lung cancer. Through a combination of in vivo tumor growth assays, immune profiling, and organoid co-culture experiments with patient-derived pancreatic tumor cells and CAFs, we demonstrate that Penetrium selectively disrupts the tumor ECM, depletes CAFs, and enhances the efficacy of immune checkpoint inhibitors and chemotherapeutic agents. Our findings support a paradigm shift in oncology: from strategies that narrowly focus on killing tumor cells to those that dismantle the malignant ecosystems in which these cells thrive.

## Results and Discussion

### Penetrium synergizes with anti-PD-1 therapy to inhibit metastatic

Penetrium, a next-generation therapeutic platform based on a repurposed niclosamide–magnesium oxide–HPMC formulation, was developed to address this critical therapeutic gap^3,26-31^. Originally derived from antiviral research, Penetrium has been reformulated to overcome the limitations of poor solubility and low tumor targeting that hampered prior attempts to apply niclosamide in oncology (Scheme 1). Engineered for selective interaction with ECM components, Penetrium exhibits enhanced stromal penetration, sustained release kinetics, and bioactivity against pathological ECM structures without harming normal tissue architecture. Importantly, its therapeutic

### progression in a 4T1 TNBC model

IVIS imaging was performed on day 27 following orthotopic 4T1 tumor implantation and 20 days of treatment (Figure 1). In the control groups (G1: vehicle; G2: anti-PD-1 alone), strong bioluminescence signals indicated a high metastatic burden in the lungs, with anti-PD-1 monotherapy showing minimal efficacy. In contrast, Penetrium monotherapy (G3: 60 mg/kg BID; G4: 120 mg/kg QD, orally) led to a dose-dependent reduction in metastatic spread, demonstrating its intrinsic anti-metastatic activity.

**Fig. 1.**
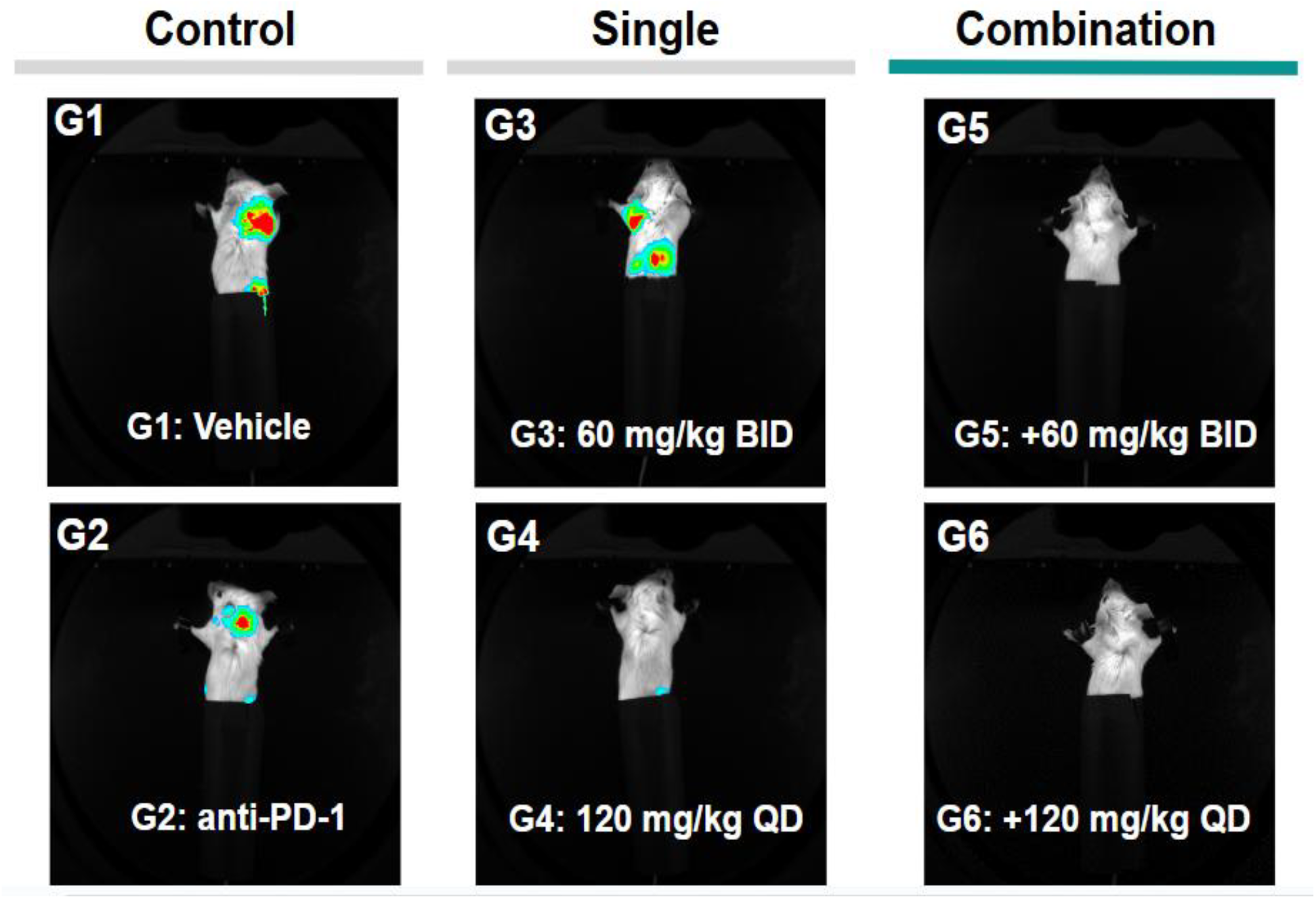
In Vivo Bioluminescence Imaging (IVIS) Reveals Therapeutic Efficacy of Penetrium and Anti-PD-1 in a 4T1 Lung Metastasis Model. IVIS imaging of lung metastases was performed on Day 27 following orthotopic injection of 4T1 breast cancer cells and 20 days after treatment initiation. Mice were treated with vehicle control (G1), anti-PD-1 monotherapy (G2), PENETRIUM monotherapy at two doses (G3: 60 mg/kg BID; G4: 120 mg/kg QD), or combination therapy with Penetrium and anti-PD-1 (G5: 60 mg/kg BID + anti-PD-1; G6: 120 mg/kg QD + anti-PD-1). Combination therapy groups (G5 and G6) showed near-complete abrogation of lung metastasis signals, demonstrating potent synergistic effects. Penetrium monotherapy (G3 and G4) reduced metastatic burden in a dose-dependent manner, while anti-PD-1 alone (G2) had minimal impact compared to vehicle (G1). Treatment administration: PENETRIUM: orally (PO) once daily or twice daily depending on the group; Anti-PD-1 antibody: 200 µg/head, intraperitoneally (IP), twice weekly (BIW)

Strikingly, combination therapy (G5 and G6) with Penetrium and anti-PD-1 antibody (200 µg/head, IP, twice weekly) resulted in near-complete suppression of lung metastases. This synergistic effect is attributed to Penetrium’s ability to remodel the tumor ECM, enabling enhanced T cell infiltration and immune recognition. Originally developed as a NIC–MgO– HPMC-based antiviral nanoplatform, Penetrium facilitates selective ECM disruption and stromal normalization, overcoming physical barriers that contribute to immune evasion and pseudo-resistance in metastatic triple-negative breast cancer.

### Penetrium Enhances Immune Cell Infiltration and Induces Localized Tumor Necrosis by Modulating the ECM Barrier

To investigate whether extracellular matrix (ECM)-mediated delivery barriers contribute to therapeutic failure in triple-negative breast cancer (TNBC), we evaluated the effects of Penetrium with and without immune checkpoint blockade in an orthotopic 4T1 tumor model. Mice were randomized into four treatment groups: (1) vehicle control, (2) anti-PD-1 monotherapy (200 µg/head, intraperitoneally, twice weekly), (3) Penetrium monotherapy (120 mg/kg, per os, once daily), and (4) combination therapy with Penetrium and anti-PD-1 antibody (Figure 2).

**Fig. 2.**
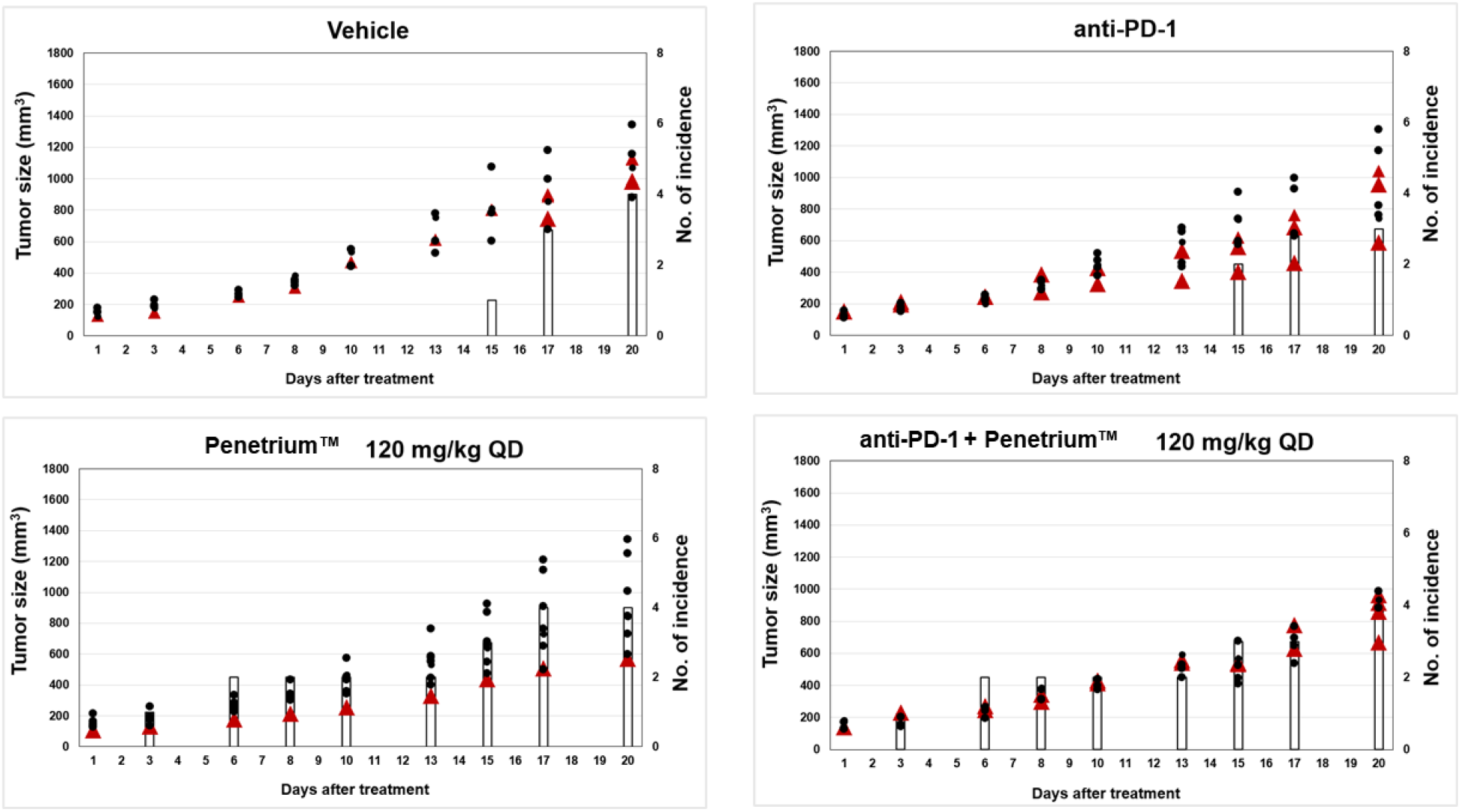
Penetrium Enhances Tumor Penetration, Immune Infiltration, and Induces Localized Necrosis in a 4T1 Breast Cancer Model. Histological and mechanistic analysis of tumors harvested from four treatment groups: (1) vehicle control, (2) anti-PD-1 monotherapy (200 µg/head, IP, twice weekly), (3) Penetrium monotherapy (120 mg/kg, orally once daily), and (4) combination therapy with anti-PD-1 and Penetrium (same doses).Localized tumor necrosis was observed in all Penetrium-treated groups (Groups 3 and 4), irrespective of the presence of immune checkpoint blockade. This necrosis was absent or minimal in the vehicle and anti-PD-1 monotherapy groups, indicating the failure of anti-PD-1 alone to elicit a therapeutic response in this ECM-rich tumor model. Mechanistically, Penetrium—a nanodrug composed of NIC–MgO–HPMC—facilitated ECM remodeling, enhancing immune cell infiltration and promoting deep intratumoral drug distribution. These findings confirm that treatment resistance in this model was primarily due to ECM-mediated delivery failure, rather than intrinsic drug insensitivity. The addition of Penetrium significantly improved immune accessibility, supporting its role as an ECM-modulating therapeutic adjuvant that converts an immune-excluded tumor into an immune-permissive one.

Histological analysis revealed pronounced localized tumor necrosis in all Penetrium-treated groups (Groups 3 and 4), while minimal or no necrosis was observed in the vehicle and anti-PD-1 monotherapy groups (Groups 1 and 2) as shown in Figure 2. These results indicate that anti-PD-1 alone was insufficient to trigger tumor cell death, consistent with the immune-excluded phenotype typical of aggressive TNBC models.

Importantly, treatment with Penetrium led to increased infiltration of immune cells into the tumor parenchyma, as confirmed by H&E and immunohistochemical staining (data not shown). This effect is attributed to Penetrium’s ability to modulate the tumor ECM. As a multifunctional nanodrug composed of niclosamide (NIC), magnesium oxide (MgO), and hydroxypropyl methylcellulose (HPMC), Penetrium is engineered for ECM penetration and remodeling. Through localized pH buffering and ionic modulation, MgO facilitates partial ECM disassembly, while HPMC stabilizes the nanoparticle and enables sustained release of NIC within the tumor microenvironment.

The enhanced necrosis and immune infiltration observed in Penetrium-treated tumors strongly suggest that therapeutic resistance in this model is not due to drug insensitivity, but rather a delivery failure caused by the dense tumor ECM. By overcoming this barrier, Penetrium reconditions the tumor stroma into an immune-permissive state, thereby rescuing the efficacy of immune checkpoint therapy when used in combination.

Together, these findings validate Penetrium as an effective ECM-modulating nanoplatform that not only restores drug delivery but also facilitates immune engagement in desmoplastic tumors like TNBC.

To evaluate the therapeutic efficacy of Penetrium in overcoming immune resistance in desmoplastic tumors, we employed an orthotopic 4T1 triple-negative breast cancer (TNBC) model, known for its aggressive growth, dense ECM, and poor response to immunotherapy. Mice bearing established 4T1 tumors were randomized into six treatment groups: (G1) vehicle control, (G2) anti-PD-1 monotherapy (200 µg/head, IP, BIW), (G3) Penetrium monotherapy at 60 mg/kg/day PO, (G4) Penetrium monotherapy at 120 mg/kg/day PO, and (G5–G6) combination therapy with anti-PD-1 and Penetrium at the same respective doses.

Following 14 consecutive days of treatment, tumor burden was quantified using **bioluminescence imaging** to measure **Total Photon Flux**. The anti-PD-1 monotherapy group (G2) failed to show any significant reduction in tumor signal compared to the vehicle group (G1), indicating that immune checkpoint blockade alone was insufficient to control tumor growth in this immune-excluded model. This aligns with previous reports describing the poor performance of PD-1/PD-L1 inhibitors in desmoplastic tumors, largely due to limited immune cell access.

In contrast, Penetrium monotherapy (G3 and G4) resulted in a dose-dependent suppression of tumor growth, with the higher dose group (G4) exhibiting a more pronounced reduction in bioluminescent signal. This finding suggests that Penetrium, originally formulated as a NIC–MgO– HPMC-based antiviral nanoplatform, possesses intrinsic antitumor activity.

**Scheme 2.**
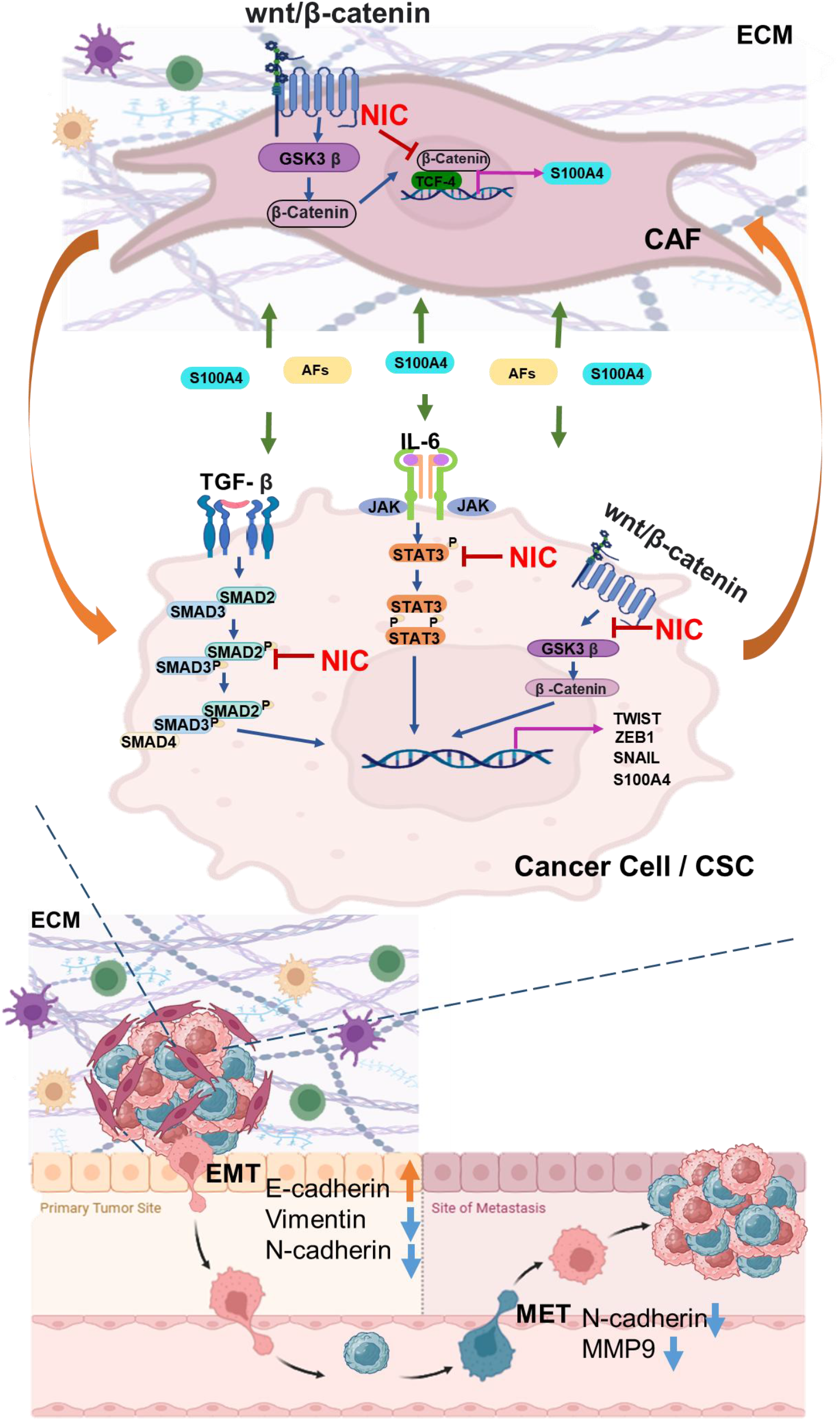
Niclosamide inhibits CAF–cancer cell crosstalk and EMT via suppression of multiple signaling pathways. Schematic illustration showing how niclosamide (NIC) disrupts key pro-tumorigenic signaling mechanisms in both cancer-associated fibroblasts (CAFs) and cancer cells. In CAFs, NIC inhibits the Wnt/β-catenin pathway, reducing S100A4 secretion and other soluble activating factors (AFs). These factors normally stimulate oncogenic STAT3 and TGF-β/SMAD signaling in adjacent cancer cells. NIC suppresses phosphorylation of STAT3 and SMAD2/3, thereby attenuating downstream transcriptional activation of EMT regulators such as TWIST, ZEB1, SNAIL, and S100A4. This leads to inhibition of epithelial-to-mesenchymal transition (EMT), reduced ECM remodeling, and decreased metastatic potential. The schematic also depicts how NIC impedes mesenchymal-to-epithelial transition (MET) and matrix metalloproteinase 9 (MMP9) activity at secondary metastatic sites.

This may be attributed to its ability to modulate the tumor microenvironment by targeting the extracellular matrix (ECM) and promoting drug diffusion into hypoxic and fibrotic tumor regions.Most notably, combination **therapy with Penetrium and anti-PD-1 (G5 and G6)** achieved a **significant and dose-dependent reduction in tumor** | **7den**, outperforming both monotherapy arms. Statistical analysis using one-way ANOVA with multiple comparisons confirmed the superiority of combination therapy:

- Compared to vehicle (G1): *p < 0*.*05 (*), p < 0.01 (**), p < 0.001 (**\*)
- Compared to anti-PD-1 alone (G2): *p < 0*.*05 (#)*

These results provide strong evidence that **Penetrium restores immune responsiveness by modulating the ECM**, thereby facilitating **T-cell infiltration** and enabling checkpoint inhibitors to engage tumor cells effectively. The ECM in solid tumors is increasingly recognized as a major contributor to **pseudo-resistance**—a phenomenon where physical barriers, rather than genetic mutations, limit therapeutic efficacy. By softening and remodeling the ECM, Penetrium disrupts these barriers, enabling immune cells and therapeutic agents to reach previously inaccessible tumor compartments.

In addition to ECM disruption, the MgO component of Penetrium may contribute to **localized pH modulation**, neutralizing the acidic tumor microenvironment and enhancing drug release kinetics. Furthermore, NIC has been shown to interfere with Wnt/β-catenin signaling and mitochondrial respiration, providing a multi-pronged therapeutic approach.

In summary, this study demonstrates that Penetrium significantly enhances the efficacy of anti-PD-1 therapy in an immune-resistant TNBC model by remodeling the ECM, facilitating immune infiltration, and enabling localized tumor killing. These findings support the development of **Penetrium as a novel immunomodulatory nanodrug platform** capable of overcoming delivery-associated resistance in solid tumors.

### Penetrium Remodels the Tumor ECM and Restores Immune Checkpoint Responsiveness in a TNBC Model

To evaluate the therapeutic efficacy of Penetrium in overcoming immune resistance in desmoplastic tumors, we employed an orthotopic 4T1 triple-negative breast cancer (TNBC) model, known for its aggressive growth, dense ECM, and poor response to immunotherapy. Mice bearing established 4T1 tumors were randomized into six treatment groups: (G1) vehicle control, (G2) anti-PD-1 monotherapy (200 µg/head, IP, BIW), (G3) Penetrium monotherapy at 60 mg/kg/day PO, (G4) Penetrium monotherapy at 120 mg/kg/day PO, and (G5–G6) combination therapy with anti-PD-1 and Penetrium at the same respective doses (Figure 3).

**Fig. 3.**
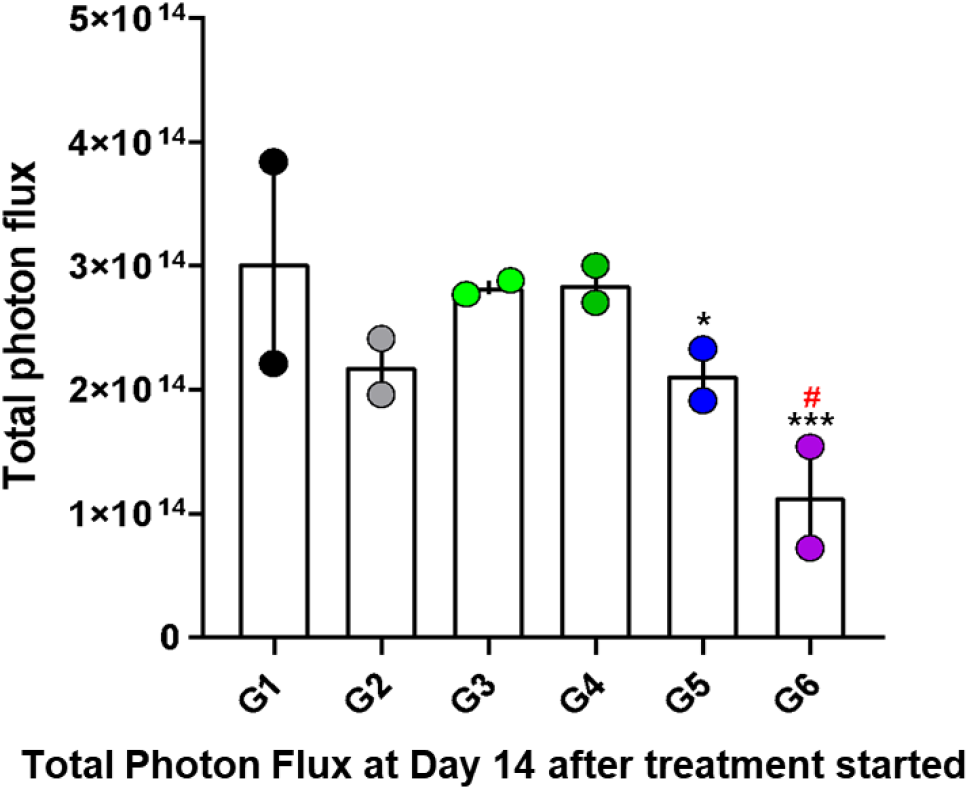
Penetrium Enhances Anti-PD-1 Efficacy and Significantly Reduces Tumor Burden After 14 Days of Treatment. Total bioluminescent signal (Total Photon Flux) was measured at Day 14 following treatment initiation in orthotopic 4T1 tumor-bearing mice. Treatment groups included: vehicle control (G1), anti-PD-1 monotherapy (G2), Penetrium monotherapy at 60 mg/kg (G3) and 120 mg/kg (G4), and combination therapy with anti-PD-1 and Penetrium at 60 mg/kg (G5) and 120 mg/kg (G6). Penetrium was administered orally once daily, and anti-PD-1 was given intraperitoneally (200 µg/head, twice weekly). Anti-PD-1 alone (G2) showed no significant reduction in tumor signal compared to vehicle, reflecting limited efficacy in this immune-excluded TNBC model. In contrast, combination therapy (G5 and G6) significantly and dose-dependently suppressed tumor burden, as evidenced by reduced Total Photon Flux, outperforming monotherapies. Statistical analysis (one-way ANOVA with multiple comparisons): Compared to G1: p < 0.05 (), p < 0.01 (), p < 0.001 (*) Compared to G2: p < 0.05 (#); These results demonstrate that Penetrium restores intratumoral T-cell infiltration and re-sensitizes tumors to immune checkpoint blockade by disrupting ECM-mediated delivery barriers.

Following 14 consecutive days of treatment, tumor burden was quantified using **bioluminescence imaging** to measure **Total Photon Flux**. The anti-PD-1 monotherapy group (G2) failed to show any significant reduction in tumor signal compared to the vehicle group (G1), indicating that immune checkpoint blockade alone was insufficient to control tumor growth in this immune-excluded model. This aligns with previous reports describing the poor performance of PD-1/PD-L1 inhibitors in desmoplastic tumors, largely due to limited immune cell access.

In contrast, **Penetrium monotherapy (G3 and G4)** resulted in a **dose-dependent suppression of tumor growth**, with the higher dose group (G4) exhibiting a more pronounced reduction in bioluminescent signal. This finding suggests that Penetrium, originally formulated as a NIC–MgO– HPMC-based antiviral nanoplatform, possesses intrinsic antitumor activity. This may be attributed to its ability to modulate the tumor microenvironment by targeting the extracellular matrix (ECM) and promoting drug diffusion into hypoxic and fibrotic tumor regions.

Most notably, **combination therapy with Penetrium and anti-PD-1 (G5 and G6)** achieved a **significant and dose-dependent reduction in tumor burden**, outperforming both monotherapy arms. Statistical analysis using one-way ANOVA with multiple comparisons confirmed the superiority of combination therapy:

- Compared to vehicle (G1): *p < 0*.*05 (*), p < 0.01 (**), p < 0.001 (**\*)
- Compared to anti-PD-1 alone (G2): *p < 0*.*05 (#)*

These results provide strong evidence that **Penetrium restores immune responsiveness by modulating the ECM**, thereby facilitating **T-cell infiltration** and enabling checkpoint inhibitors to engage tumor cells effectively. The ECM in solid tumors is increasingly recognized as a major contributor to **pseudo-resistance**—a phenomenon where physical barriers, rather than genetic mutations, limit therapeutic efficacy. By softening and remodeling the ECM, Penetrium disrupts these barriers, enabling immune cells and therapeutic agents to reach previously inaccessible tumor compartments.

NIC has been shown to interfere with Wnt/β-catenin signaling and mitochondrial respiration, providing a multi-pronged therapeutic approach. In summary, this study demonstrates that Penetrium significantly enhances the efficacy of anti-PD-1 therapy in an immune-resistant TNBC model by remodeling the ECM, facilitating immune infiltration, and enabling localized tumor killing. These findings support the development of **Penetrium as a novel immunomodulatory nanodrug platform** capable of overcoming delivery-associated resistance in solid tumors.

To investigate immune-related mechanisms underlying the therapeutic response, we performed quantitative RT-PCR on tumor and lung tissue from mice treated with vehicle, anti–PD-1, HAB-SON01, or their combination (Figure S0). Tumor analysis revealed a trend toward increased **CD8**+ T cell signatures in the combination group (G11), although variability was observed. Expression of **Ly6e**, associated with inflammatory monocytes and early neutrophil infiltration, was significantly elevated in G11 (*p* < 0.05), suggesting enhanced immune cell recruitment. Furthermore, expression of **F4/80**, a marker for tumor-associated macrophages (TAMs), was also elevated in the G11 group, indicating active immune engagement. Importantly, **TGF-β**, a hallmark immunosuppressive cytokine, was robustly downregulated across all treatment arms, with the greatest reduction in the combination group. This indicates attenuation of tumor-mediated immune suppression.

Systemic immunotoxicity was assessed via **Arg1** expression in lung tissue. No significant differences were observed across groups, suggesting that HAB-SON01, even in combination, does not provoke peripheral immunosuppression or M2-skewed responses.

These findings support the hypothesis that **HAB-SON01 acts as an immune-potentiating agent** when combined with checkpoint inhibition, enhancing antitumor immunity without off-target immune activation.

### Dual Targeting of ECM and Angiogenesis by Penetrium and Bevacizumab Synergistically Suppresses Metastasis

To evaluate whether simultaneous modulation of the tumor extracellular matrix (ECM) and inhibition of angiogenesis could effectively suppress metastatic progression, we employed the LL/2 lung carcinoma xenograft model. Mice were treated with bevacizumab (BVZ; VEGF inhibitor, 5 mg/kg, IP, BIW) alone or in combination with escalating doses of Penetrium (50, 100, or 150 mg/kg, orally, QD). Penetrium, a NIC–MgO– HPMC nanodrug, was hypothesized to remodel the ECM and facilitate deeper penetration of immune or antiangiogenic agents (Figure 4).

**Fig. 4.**
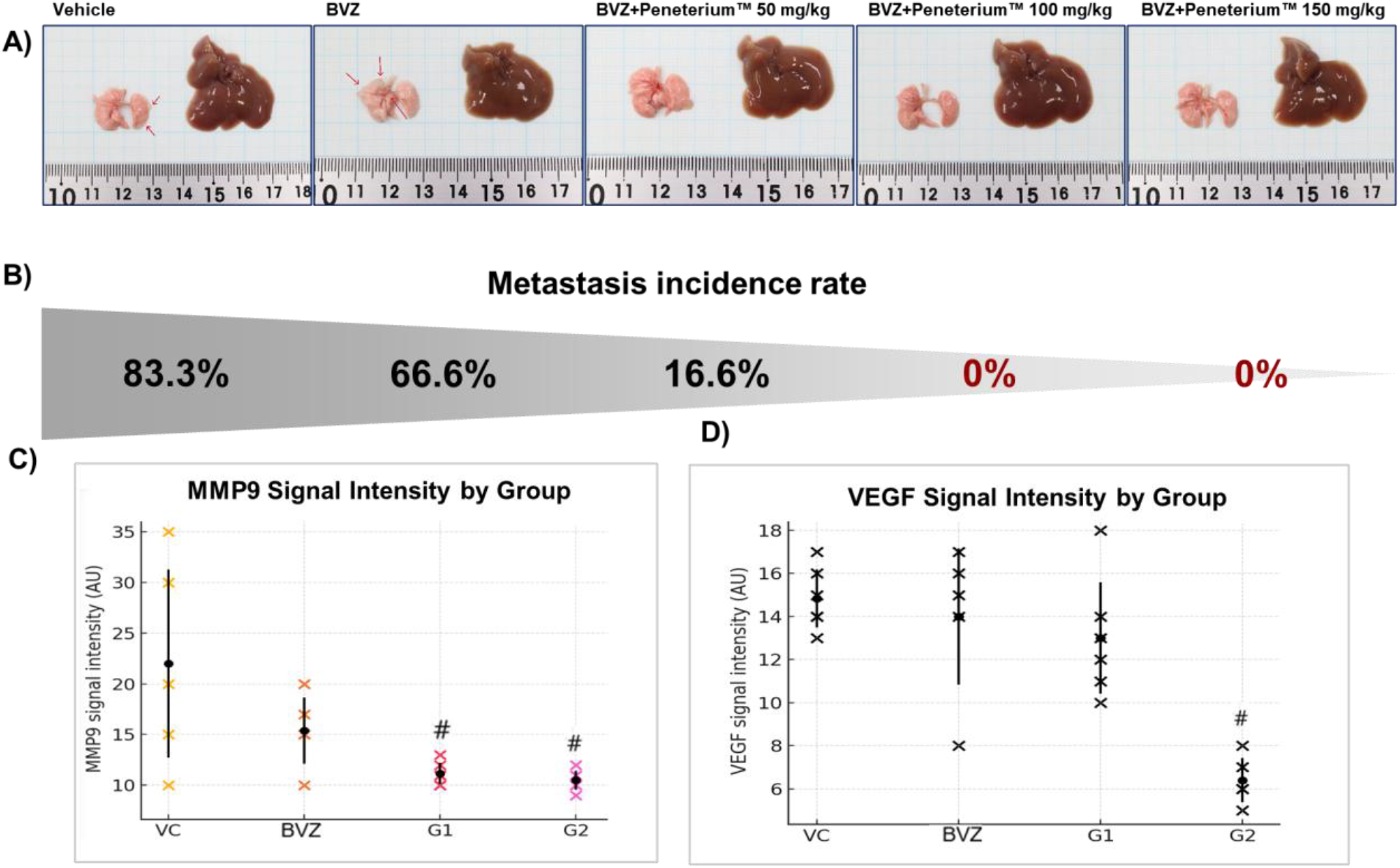
Penetrium Synergizes with Bevacizumab to Inhibit Angiogenesis and Suppress Metastasis through Dual Targeting of ECM and VEGF Signaling: In vivo therapeutic efficacy of Penetrium and bevacizumab (BVZ) combination therapy was assessed in the LL/2 subcutaneous xenograft mouse model, focusing on anti-angiogenic response, metastatic suppression, and tumor microenvironment remodeling. (A, B) Representative images show vascular density in tumor tissues across different treatment groups. The upper panels compare: Low/partial anti-angiogenic response in vehicle and BVZ monotherapy groups, Versus marked suppression of angiogenesis in combination groups treated with BVZ + Penetrium at escalating doses (50 mg/kg, 100 mg/kg, and 150 mg/kg, administered orally, QD). Bevacizumab was given intraperitoneally (5 mg/kg, BIW). Vascular rarefaction and architectural disruption were most pronounced in the high-dose Penetrium combination groups; (B) Quantification of metastatic incidence (% metastatic rate) shows a dose-dependent reduction in metastasis with combination therapy, significantly outperforming both vehicle and BVZ monotherapy groups; (C, D) Mechanistic biomarker analysis shows significant downregulation of: MMP-9 (C), a matrix metalloproteinase associated with ECM degradation and metastatic invasion, VEGF (D), a pro-angiogenic factor directly targeted by BVZ.; Notably, Penetrium monotherapy moderately reduced MMP-9 levels, while its combination with BVZ showed additive or synergistic suppression of both MMP-9 and VEGF, suggesting dual microenvironmental targeting— of both the stromal matrix and vasculature. Together, these findings highlight the synergistic anti-metastatic effect of Penetrium and BVZ, and demonstrate that dual targeting of ECM remodeling and angiogenesis provides a rational and potent strategy to inhibit tumor progression and distant metastasis in highly invasive cancers.

Macroscopic and histological examination of tumor vasculature (Figure 4A, B upper panels) revealed that the vehicle and BVZ monotherapy groups retained extensive vascular networks, indicating that BVZ alone only partially inhibited angiogenesis in this aggressive tumor model. In contrast, the BVZ + Penetrium groups showed a dose-dependent reduction in microvascular density, with the 100 mg/kg and 150 mg/kg Penetrium groups displaying nearly collapsed or poorly perfused vasculature, suggestive of vascular normalization or regression.

This anti-angiogenic effect correlated strongly with reduced metastatic incidence, as shown in Figure 4B (lower panel). Quantitative analysis demonstrated that the percentage of mice with metastatic lesions was significantly lower in all combination therapy groups, especially at higher Penetrium doses. These data suggest that Penetrium enhances the anti-metastatic efficacy of BVZ, likely by disrupting ECM barriers and improving the functional delivery of antiangiogenic agents to distal tumor niches.

To explore underlying mechanisms, we assessed the expression of MMP-9 and VEGF, key molecular drivers of matrix remodeling and neovascularization. As shown in Figures 4C and D, Penetrium monotherapy moderately suppressed MMP-9, while BVZ alone inhibited VEGF but had minimal effect on MMP-9. Strikingly, the combination therapy synergistically downregulated both MMP-9 and VEGF, suggesting a dual blockade of ECM turnover and angiogenic signaling.

These findings support the hypothesis that co-targeting the tumor microenvironment via ECM modulation (Penetrium) and angiogenesis inhibition (BVZ) is a rational strategy for metastatic suppression. Penetrium appears to play a dual role—facilitating drug access by disrupting dense ECM structures, and directly modulating the tumor microenvironment by suppressing MMP activity. When combined with VEGF blockade, this leads to a collapse of pro-metastatic signaling pathways and a significant reduction in metastatic spread.

### Penetrium Synergizes with Paclitaxel to Suppress Metastasis and Reverse EMT in the EO771 Syngeneic Breast Cancer Model

To evaluate the therapeutic potential of Penetrium in improving the efficacy of chemotherapy against metastatic breast cancer, we employed the EO771 syngeneic mouse model—an immunocompetent system that closely mimics spontaneous metastasis. Tumors were established by intravenous injection of EO771 cells, and mice were treated for 14 days with various regimens: vehicle (G1), paclitaxel (PTX, 2 mg/kg EOD, G2), Penetrium monotherapy (200 mg/kg/day, G3), or PTX in combination with Penetrium at 100 mg/kg/day (G4) or 200 mg/kg/day (G5).

Quantitative assessment of lung metastasis, expressed as the **relative tumor area over total lung surface**, revealed a **significant and dose-dependent suppression of metastatic burden** in groups receiving Penetrium (Figure 5A). Monotherapy with PTX (G2) showed modest efficacy, while combination therapy (G4, G5) achieved robust anti-metastatic effects, with the high-dose group (G5) showing the most significant improvement. Statistical analysis demonstrated strong significance:

**Fig. 5.**
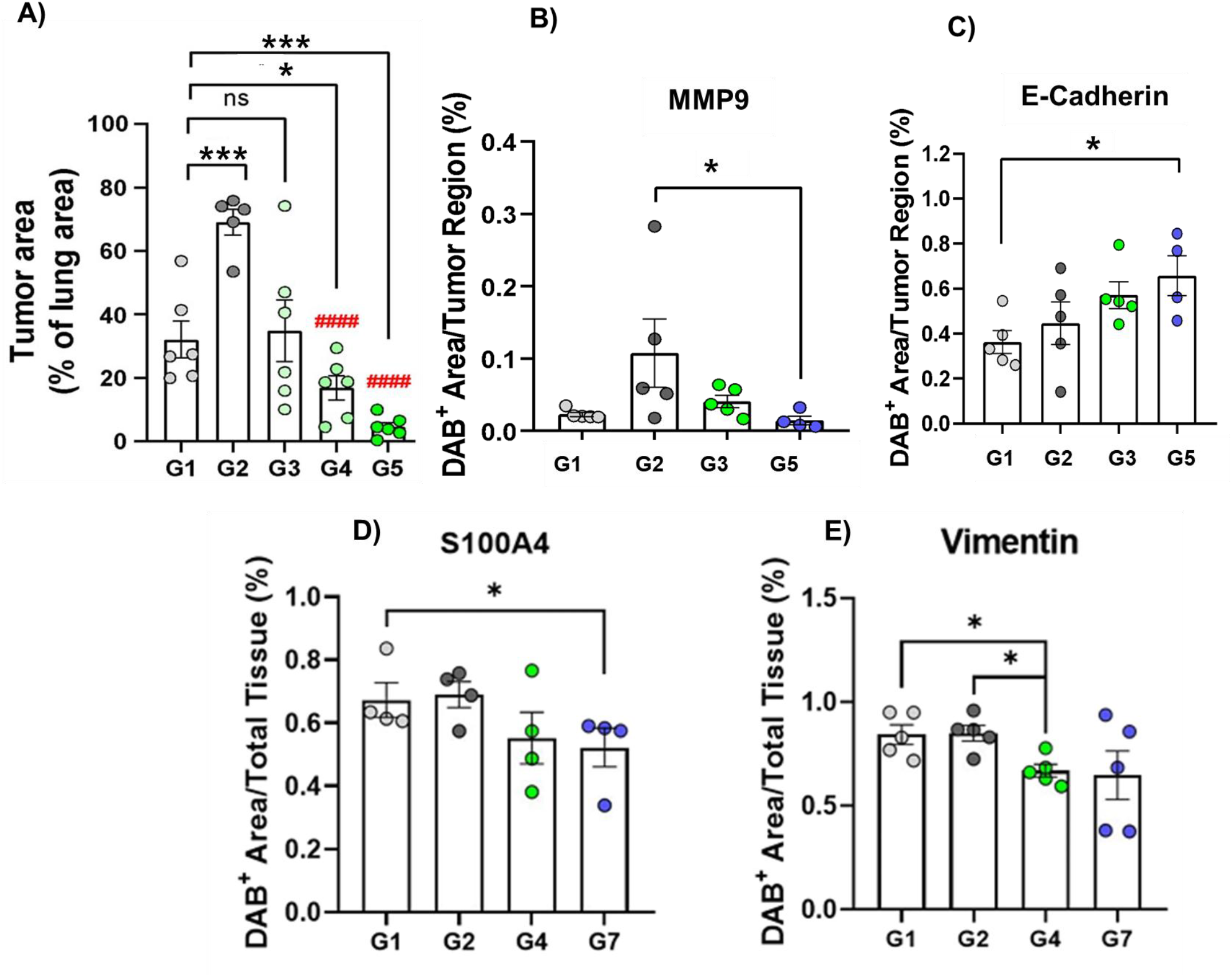
Penetrium Enhances Paclitaxel Efficacy, Suppresses Metastasis, and Modulates EMT Markers in EO771 Syngeneic Model. In vivo evaluation of Penetrium in combination with paclitaxel (PTX) was performed using the EO771 syngeneic breast cancer model via intravenous tumor cell injection, designed to mimic spontaneous lung metastasis. Mice were treated with: G1: Vehicle control, G2: PTX monotherapy (2 mg/kg, every other day, IV), G3: Penetrium monotherapy (200 mg/kg/day, oral), G4: Combination therapy with PTX + Penetrium 100 mg/kg/day, G5: Combination therapy with PTX + Penetrium 200 mg/kg/day; (A) Quantification of lung metastases, based on the relative tumor area to total lung surface, showed a dose-dependent reduction in metastatic burden in Penetrium-treated groups. Notably, combination therapy in G5 (PTX + Penetrium 200 mg/kg/day) exhibited the most significant suppression of lung metastasis; (Statistical analysis (one-way ANOVA, n = 6 per group, mean ± SEM): Compared to G1: p < 0.05 (), p < (), p < 0.001 (), p < 0.0001, Compared to G2: #p < 0.05, ####p < 0.0001; (B) Immunohistochemical analysis revealed that Penetrium reduces the expression of MMP-9, a key ECM-degrading enzyme associated with metastatic invasion and tumor cell dissemination; (C) Furthermore, Penetrium restored epithelial characteristics, evidenced by the upregulation of E-cadherin, a hallmark of epithelial phenotype. Combination therapy (G5) demonstrated the strongest re-expression of E-cadherin, suggesting reversal of epithelial–mesenchymal transition (EMT) and reduction in metastatic potential; Collectively, these results confirm that Penetrium enhances the anti-metastatic efficacy of paclitaxel while also exerting microenvironmental and phenotypic remodeling effects—via MMP-9 suppression and EMT reversal—providing a robust rationale for its use in metastatic breast cancer therapy. **(D–E)** Additional EMT-associated markers—S100A4 and vimentin—were significantly downregulated in Penetrium-treated groups (G4, G5, G7), confirming the suppression of mesenchymal traits. Overall, these results suggest that Penetrium synergizes with paclitaxel to inhibit metastatic progression by suppressing EMT and ECM remodeling, offering a promising combinatorial strategy for advanced breast cancer treatment.

- Compared to vehicle (G1): *p < 0*.*05 (*), **p < 0.01 (**), ***p < 0*.*001 (***), **p < 0.0001 (**)
- Compared to PTX (G2): #p < 0.05, ####p < 0.0001

To further elucidate the mechanistic basis of this effect, we evaluated the expression of **matrix metalloproteinase-9 (MMP-9)** and **E-cadherin** via immunohistochemistry. MMP-9, a key ECM-degrading enzyme that facilitates cancer cell invasion and metastasis, was markedly suppressed by Penetrium treatment (Figure 5B). In the combination group (G5), MMP-9 expression was significantly downregulated, correlating with reduced ECM disruption and metastatic spread.

Importantly, Penetrium also restored **E-cadherin expression** (Figure 5C), a hallmark of epithelial identity that is commonly lost during epithelial– mesenchymal transition (EMT). The re-expression of E-cadherin was most evident in the combination group (G5), suggesting that **Penetrium not only inhibits ECM remodeling but also reverses EMT**, thereby attenuating metastatic potential at both structural and phenotypic levels.

These findings confirm that Penetrium enhances paclitaxel’s therapeutic efficacy by **modulating the tumor microenvironment**, suppressing ECM-associated invasion (via MMP-9 inhibition), and reprogramming metastatic cells toward an epithelial state (via E-cadherin upregulation). The **dose flexibility** of Penetrium in combination therapy underscores its clinical potential as a synergistic co-therapeutic that both **improves primary tumor control** and **prevents systemic metastasis** in aggressive breast cancers.

**A graphical summary of the proposed mechanism is presented in Scheme 2**, illustrating how Penetrium (via its active component, niclosamide) inhibits CAF-mediated signaling, suppresses key pathways such as STAT3, SMAD, and Wnt/β-catenin, and consequently reprograms cancer cells toward an epithelial phenotype by restoring E-cadherin and downregulating MMP-9 and S100A4 expression.

### Penetrium Enhances Stromal Suppression in Pancreatic Tumor Organoid Co-Culture Organoids

To investigate the therapeutic advantage of nanohybridization, we compared the cytotoxic performance of native niclosamide with its nanoengineered analog, Penetrium, in a patient-derived pancreatic tumor model incorporating both organoids and stromal CAFs (Figure 6). Pancreatic cancer organoids (PCOs) derived from hPC 21-016 were cultured alone or in combination with patient-matched CAFs to recapitulate a desmoplastic microenvironment. Treatments were performed using increasing concentrations of either niclosamide or Penetrium (0.25, 0.5, 1, and 2 µM), and relative viability was assessed after 5 days. Area under the curve (AUC) calculations were used to quantify cumulative cytotoxicity for each condition.

**Figure 6.**
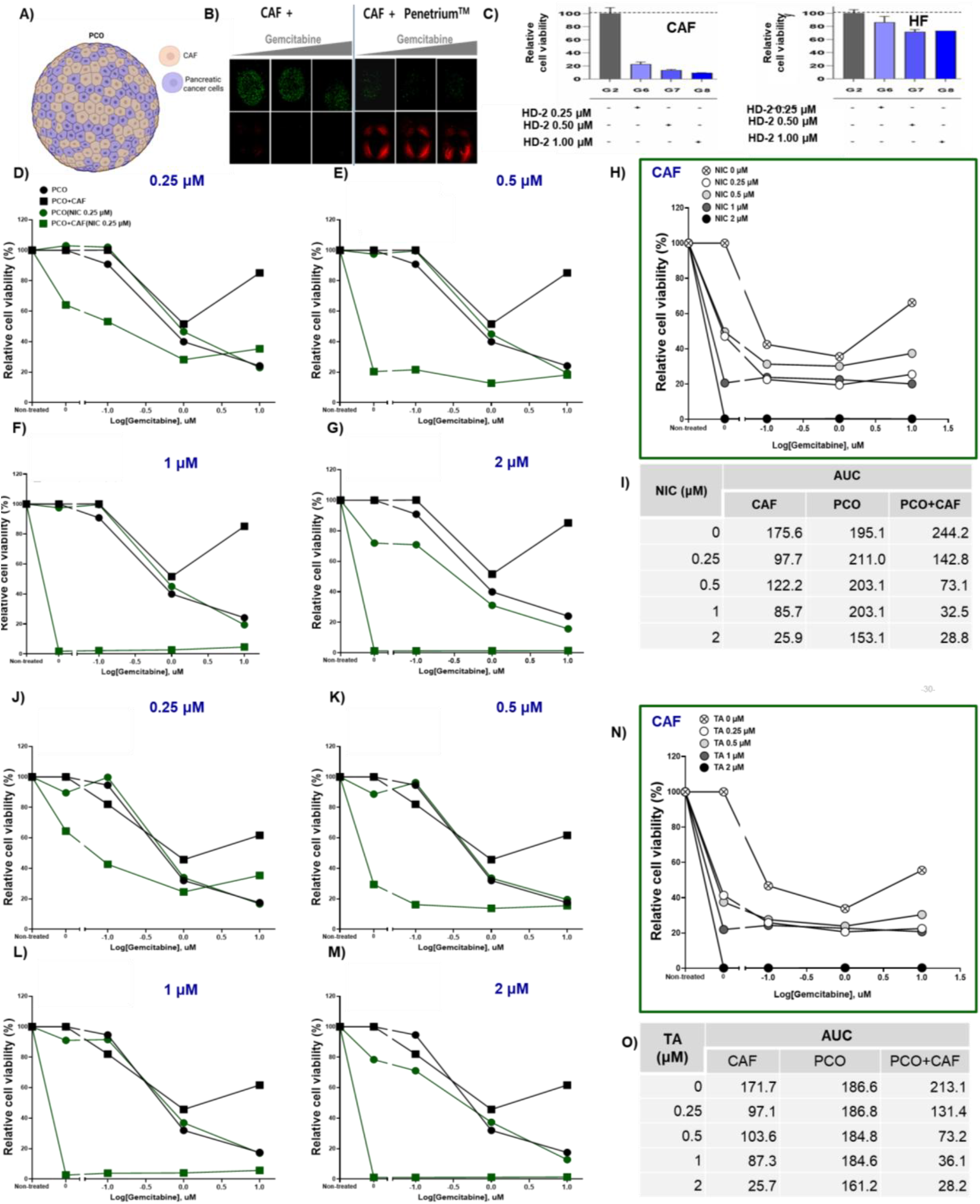
Selective cytotoxicity of HD-2 towards cancer-associated fibroblasts (CAFs) over normal human fibroblasts. **(A)**Represents PCO+CAF organoid culture model; (B) Live/dead fluorescence staining visualizes the cell fate outcome. Green fluorescence (Calcein AM) indicates live cells; red fluorescence (Ethidium Homodimer-1) marks dead cells. The data highlight HD-2’s preferential cytotoxicity against CAFs while sparing normal fibroblasts, (C) Comparative analysis in normal human fibroblasts (HFs) shows significantly less cytotoxicity at equivalent HD-2 doses, confirming selective targeting of CAFs. validating its tumor stroma– targeting potential. (Scale bar = 10µm); Representative images shown; N=3 independent experiments) and Relative viability of CAFs after 24 h exposure to increasing concentrations of HD-2 (0.25, 0.5, and 1 μM). A dose-dependent decrease in CAF viability is observed, with G8 (1 μM) showing the strongest effect. **(D-I and J-O) Comparative cytotoxic evaluation of niclosamide and Penetrium in patient-derived pancreatic cancer organoid (PCO) and cancer-associated fibroblast (CAF) co-culture models**. Dose–response analysis was conducted over 5 days in mono-and co-culture settings using niclosamide (top panels) and Penetrium (bottom panels) across four concentrations (0.25–2 µM). Relative cell viability curves are shown for CAF, PCO, and PCO+CAF models. Dot plots illustrate area under the curve (AUC) values representing cumulative cytotoxicity, and tables summarize the quantitative AUC data. Both formulations exhibited dose-dependent suppression of viability, with markedly enhanced cytotoxicity in the co-culture setting. Penetrium showed greater CAF-specific suppression at intermediate doses, suggesting enhanced stromal penetration and therapeutic synergy in desmoplastic tumor contexts (n=3, * p <0.05).

Both niclosamide and Penetrium elicited strong, dose-dependent reductions in cell viability. In the co-culture condition, where PCOs and CAFs were grown together, the AUC dropped significantly—from 244.2 (untreated) to 28.8 at 2 µM for niclosamide, and similarly from 213.1 to 28.2 for Penetrium. These reductions were more pronounced than those seen in either monoculture, suggesting synergistic efficacy when targeting both tumor and stromal compartments simultaneously.

Interestingly, Penetrium demonstrated enhanced suppression of CAF viability at intermediate doses (0.5–1 µM), with lower AUC values than those observed for native niclosamide. This observation supports the hypothesis that nanohybridization improves drug delivery and cellular engagement within the stromal compartment, possibly via increased uptake or sustained exposure through the MgO–HPMC matrix. Since CAFs are key contributors to ECM remodeling and therapeutic resistance, their preferential elimination at lower drug concentrations may facilitate greater tumor vulnerability and deeper therapeutic penetration.

These findings demonstrate the capacity of Penetrium to target not only malignant epithelial cells but also the supportive fibroblastic niche that enables tumor progression. This dual-compartment action provides a critical advantage in the treatment of desmoplastic cancers such as PDAC, where stromal shielding remains a central obstacle to effective drug delivery. The data also validate the utility of nanohybrid drug design as a strategic means of optimizing biodistribution and achieving spatially coordinated therapeutic effects across tumor microenvironmental compartments.

To further elucidate the therapeutic impact of nanohybridization, we evaluated the cumulative cytotoxicity of Penetrium versus native niclosamide in the same organoid-CAF co-culture model over a 5-day exposure window (Figure S1). Both formulations produced dose-dependent decreases in viability, but Penetrium consistently exhibited stronger effects in the stromal co-culture condition. Notably, at intermediate concentrations (0.5–1 µM), Penetrium induced lower AUC values compared to free niclosamide, indicating enhanced CAF suppression. This suggests that the MgO–HPMC nanomatrix may provide sustained drug exposure or improved cellular uptake, particularly within the stromal compartment.

Given the central role of CAFs in matrix remodeling and drug resistance, their preferential targeting by Penetrium may render the surrounding tumor tissue more permeable and susceptible to treatment. This dual-compartment action—simultaneously suppressing malignant epithelial cells and their protective fibroblastic niche—underscores the translational promise of ECM-modulating nanotherapeutics. It also validates nanohybrid drug design as a means to achieve spatially coordinated therapeutic effects across distinct compartments of the tumor microenvironment.

To validate the dual-compartment anticancer activity of Penetrium and benchmark it against native niclosamide, we tested both compounds in an independent patient-derived pancreatic cancer organoid (PCO) and cancer-associated fibroblast (CAF) co-culture model (hPC 20-001). Co-cultures and monocultures were treated for 5 days with increasing concentrations of niclosamide or Penetrium (0.25–2 µM), and relative cell viability was assessed. Cumulative cytotoxicity was quantified via AUC calculations.

Both formulations induced a clear dose-dependent reduction in cell viability across all culture conditions. In the co-culture setting, AUC dropped from 240.0 (untreated) to 46.6 at 2 µM niclosamide, and from 198.1 to 28.0 with Penetrium, indicating potent tumor–stroma co-elimination. Notably, Penetrium consistently demonstrated **greater CAF suppression at intermediate concentrations** (0.5–1 µM), with CAF AUCs of 216.2 and 91.0, compared to 207.7 and 90.5 with niclosamide, respectively.

At higher doses (2 µM), both compounds achieved near-complete ablation of CAFs, but Penetrium produced greater co-culture suppression despite slightly higher PCO viability—suggesting an improved stromal modulation effect that enhances overall efficacy. These differences are likely due to the nanohybrid structure of Penetrium, which may offer improved penetration, stability, and sustained release properties that increase drug accumulation within the stromal compartment.

This second patient-derived model reinforces the findings from the hPC 21-016 study, confirming that nanohybridization of niclosamide enhances dual-compartment cytotoxicity in desmoplastic tumor systems (Figure S2). The improved performance of Penetrium, particularly at clinically relevant concentrations, highlights its translational promise as a **stromal-sensitizing anticancer therapeutic** with potential to overcome ECM-mediated resistance barriers that are characteristic of pancreatic ductal adenocarcinoma. These results support the rationale for advancing Penetrium into preclinical efficacy studies in **orthotopic, stroma-rich in vivo models**, with the ultimate aim of developing **multi-compartment-targeted treatments for otherwise drug-refractory solid tumors**.

To evaluate the effect of Penetrium on the tumor-stroma axis, we designed a co-culture model comprising patient-derived pancreatic cancer organoids (PCOs) and matched cancer-associated fibroblasts (CAFs), mimicking the desmoplastic microenvironment of PDAC. Organoids and CAFs were cultured individually (monoculture) and in combination (PCO+CAF) and treated with escalating doses of either gemcitabine, niclosamide, or Penetrium. Viability was assessed after 5 days, and the cumulative drug effect was quantified by calculating area under the curve (AUC) values across the dose-response curves.

As shown in **Figure S3** gemcitabine exhibited a reduction in PCO viability (AUC 140.7), but its effect was blunted in CAFs (AUC 217.2) and remained high in the co-culture condition (AUC 180.4), suggesting a stromal-mediated protective effect. In contrast, niclosamide significantly reduced viability in both CAFs (AUC 59.9) and the co-culture (AUC 55.9), demonstrating its dual-compartment cytotoxicity. Most notably, Penetrium achieved comparable suppression of co-culture viability (AUC 54.8) but exhibited enhanced CAF-specific toxicity (AUC 89.4) compared to niclosamide. Despite slightly higher AUC in CAFs, Penetrium retained a lower AUC in the co-culture than gemcitabine, indicating its superior ability to overcome stromal interference.

These results validate the hypothesis that nanoformulation of niclosamide improves spatial pharmacodynamics by enhancing bioavailability and targeting both malignant and stromal cells within the tumor microenvironment. The ability of Penetrium to penetrate and disrupt the fibroblastic niche positions it as a promising candidate for treating desmoplastic tumors where conventional therapies often fail to reach their cellular targets.

### Selective Stromal Vulnerability Revealed by NIC-Gemcitabine Response Profiling

To further delineate the impact of niclosamide on distinct tumor microenvironmental compartments, we investigated how pre-treatment with increasing doses of niclosamide affected the cytotoxic response to gemcitabine in isolated and combined PCO–CAF cultures. As shown in **Figure S4**, treatment with gemcitabine following niclosamide exposure revealed a marked shift in drug sensitivity profiles.

In organoid-only systems, gemcitabine efficacy remained relatively stable across NIC concentrations, with IC50 values ranging from 0.8 to 0.36 µM. However, CAF-only cultures exhibited a dramatic loss in viability when treated with 1–2 µM of niclosamide, as evidenced by the collapse in total peak area (from 345.2 to 3.71) and a corresponding IC50 surge (from ∼1.0 to 7033 µM). In the co-culture condition, gemcitabine IC50 remained suppressed (1.078 µM at NIC 2 µM), indicating that tumor drug responsiveness improved when stromal elements were selectively eliminated. This concentration-dependent stromal toxicity suggests that niclosamide acts not only as a tumor cytotoxin but also as a stroma-modulating agent that neutralizes CAF-mediated chemoresistance. These findings align with our prior observations of reduced AUCs in co-culture and support the development of Penetrium for its dual-compartment targeting properties. To evaluate the extent of stromal interference on standard chemotherapy, we investigated the efficacy of gemcitabine at three concentrations (0.1, 1, and 10 µM) across PCO monoculture, CAF monoculture, and PCO+CAF co-culture systems. As shown in **Figure S5** gemcitabine showed strong cytotoxicity in PCOs at all concentrations (AUC 284 → 22.59; IC50 0.098 → 3.1 µM). However, in CAF-only cultures, AUC values remained elevated (270.2–207.2), with IC50 values consistently >0.75 µM.

More strikingly, the co-culture model showed minimal changes in AUC (from 147.8 to 146.7) despite a 100-fold increase in gemcitabine concentration, and IC50 values remained drastically elevated (1,943,301 at 0.1 µM, 15,875 at 1 µM), reflecting **pseudo-resistance induced by the stromal component**. Even at 10 µM, IC50 dropped to only 1.05 µM in co-culture, while CAFs retained resistance.

These findings clearly demonstrate that **CAF presence significantly blunts gemcitabine efficacy** even at high doses and further justify the development of dual compartment targeting agents like Penetrium, which are capable of selectively suppressing the stromal shield.

To validate this phenomenon further, we tested **niclosamide (referred to here as HD-2)** for its selective cytotoxicity toward CAFs. As shown in **Figure 6A-C**, niclosamide induced a strong, dose-dependent reduction in CAF viability, while sparing normal human fibroblasts. Live/dead staining confirmed this selectivity, revealing extensive cell death in CAFs at 1 μM, whereas normal fibroblasts remained largely viable. These findings provide direct evidence that **niclosamide preferentially targets the stromal compartment**, particularly CAFs, which are known contributors to therapeutic resistance. This supports our mechanistic hypothesis that **Penetrium—a niclosamide-based nanohybrid—acts as a dual-compartment therapeutic**: directly cytotoxic to tumor cells while simultaneously dismantling the stromal shield that drives chemoresistance.

## Conclusions

This study establishes Penetrium, a NIC–MgO–HPMC-based multifunctional nanodrug, as a powerful therapeutic platform that addresses a fundamental barrier in solid tumor treatment—extracellular matrix (ECM)-mediated pseudo-resistance. Across multiple preclinical models of aggressive and immune-resistant cancers, including orthotopic 4T1 triple-negative breast cancer (TNBC), EO771 syngeneic breast cancer, and LL/2 lung carcinoma, Penetrium consistently demonstrated robust anti-metastatic efficacy and synergistic enhancement of standard therapies. In the context of immune checkpoint blockade, Penetrium enabled restoration of T cell infiltration, reversed immune exclusion, and rescued anti-PD-1 efficacy—achieving near-complete suppression of lung metastasis in combination therapy. This was mechanistically attributed to Penetrium’s ability to remodel the ECM, allowing immune cells to access tumor niches previously inaccessible due to stromal density and fibrosis. In chemotherapeutic and antiangiogenic contexts, Penetrium significantly amplified the effects of paclitaxel and bevacizumab, respectively, by modulating the tumor microenvironment at multiple levels. This included suppression of MMP-9, restoration of E-cadherin expression, and inhibition of VEGF-mediated angiogenesis, indicating reversal of epithelial– mesenchymal transition (EMT) and blockade of metastatic signaling cascades. Importantly, Penetrium exhibited dose-dependent activity, offering flexibility for therapeutic customization without compromising efficacy. Its bioinspired composition and dual-action profile—combining localized pH buffering, controlled NIC release, and matrix remodeling— provide a multi-pronged approach to overcoming drug delivery barriers and enhancing treatment responses. Together, these findings position Penetrium as a next-generation nanotherapeutic capable of transforming the treatment paradigm in solid tumors by converting immune-excluded, desmoplastic, and therapy-resistant tumors into immune-permissive and drug-accessible landscapes. Its integration into existing oncologic regimens holds considerable promise for improving outcomes in patients with metastatic and refractory cancers.

## Methods

### Experimental Section

#### Penetrium (CP-COV03) Nanohybrid Synthesis via Solid–Liquid Interface Activation Method

Penetrium (also referred to as CP-COV03) was synthesized using a nanohybridization strategy involving a solid–liquid interface activation technique, optimized for clinical-grade production. The formulation was manufactured in a GMP-certified facility (YOOYOUNG Pharm., South Korea) under standardized quality protocols. In brief, niclosamide (2,400 g), magnesium oxide (MgO; 1,680 g), and hydroxypropyl methylcellulose (HPMC; 600 g) were precisely weighed and premixed manually. From this bulk mixture, 195 g was subjected to initial homogenization using a planetary ball mill (Retsch PM100) at 170 rpm for 10 minutes. A solvent solution containing 419.95 g of 95% ethanol and 277.76 g of distilled water was prepared, and 20.8 g of the powdered mixture was added to this solvent phase. The resulting slurry underwent five sequential milling cycles (2 minutes each) at 170 rpm, followed by four extended milling cycles of 10 minutes each under the same rotational conditions. The milled nanohybrid suspension was then dried using a vacuum dryer set at 45 °C for at least 1 hour to yield the final nanohybrid powder. This dried nanocomposite material was then encapsulated into hard gelatin capsules, each containing 50 mg of niclosamide. The final product, Penetrium, is a clinically scalable and pharmaceutically stable nanohybrid formulation designed to enhance drug bioavailability and stromal penetration.

#### Animal model

All procedures involving animals were conducted in accordance with ethical guidelines and were approved by the relevant Institutional Animal Care and Use Committee (IACUC). C57BL/6 (male, 6–8 weeks old) and BALB/c (male and female, 6–8 weeks old) mice were obtained from OrientBio (Seoul, Korea). Mice were housed in a controlled environment with free access to food and water. Upon reaching a tumor volume of 50–100 mm^3^, animals were randomly assigned to experimental groups and received PENETRIUM or combination treatments as outlined in Table 1.

**Table 1.** Treatment regime.

|  |  |
| --- | --- |
| 4T1 orthotopic mouse model |  |
| Group1 | Vehicle |
| Group2 | Anti-PD-1 (200 ug/head, IP, BIW) |
| Group3 | Penetrium 120 mg/kg, PO, QD |
| Group4 | Anti-PD-1 → Anti-PD-1 + Penetrium 120 mg/kg, PO, QD |
| Group5 | Anti-PD-1 + Penetrium 120 mg/kg, PO, QD |
| EO771 mouse model |  |
| Group1 | Vehicle |
| Group2 | Paclitaxel (2 mg/kg, IV, EOD) |
| Group3 | Penetrium 200 mg/kg, PO, QD |
| Group4 | Paclitaxel + 100 mg/kg, PO, QD |
| Group5 | Paclitaxel + 200 mg/kg, PO, QD |
| LL/2 lung metastasis model |  |
| Group1 | Vehicle |
| Group2 | Bevacizumab (5 mg/kg, IP, BIW) |
| Group3 | Penetrium 50, 100, or 150 mg/kg, PO, QD |
| Group4 | Bevacizumab + Penetrium 50, 100, or 150 mg/kg, PO, QD |

To generate the orthotopic triple-negative breast cancer (TNBC) model, 2 × 105 4T1-Flu cells were injected into the fourth mammary fat pad of female BALB/c mice. Tumor progression and metastatic spread to the lungs were monitored using bioluminescent imaging (IVIS Lumina II, PerkinElmer, Waltham, MA, USA). D-luciferin potassium salt (150 mg/kg, i.p.) was administered prior to imaging, and bioluminescence was recorded 3 minutes post-injection at predetermined time points. Tumor burden was assessed by quantifying total photon flux.

In the EO771 lung metastasis model, C57BL/6 mice were intravenously injected with 5 × 105 EO771 cells. Therapeutic interventions were initiated three days after cell inoculation.

For the LL/2 lung carcinoma xenograft model, C57BL/6 mice were subcutaneously implanted with 5 × 105 LL/2 cells. Metastatic frequency was evaluated using hematoxylin and eosin (H&E) staining, while protein expression of VEGF and MMP-9 was determined through immunohistochemistry (IHC).

#### Immunohistochemistry

Tissue samples were fixed in 4% paraformaldehyde (Biosolution Co., Ltd., Seoul, Korea) for 24 hours (tissues) or 30 minutes (organoids), followed by paraffin embedding. Sections (4 µm thick) were processed through xylene and a graded ethanol series for deparaffinization and rehydration, then stained with H&E or Masson’s trichrome (Dako, Carpinteria, CA, USA). For IHC, antigen retrieval was performed in sodium citrate buffer (10 mM, pH 6.0, with 0.05% Tween-20) at 95°C for 1 hour. Endogenous peroxidase activity was blocked using 3% hydrogen peroxide in methanol for 10 minutes. Sections were then incubated with 1% bovine serum albumin (BSA; GenDEPOT, TX, USA) in PBS for blocking, followed by overnight incubation at 4°C with primary antibodies targeting MMP-9 (ab22402, Abcam, 1:100) and VEGF (catalog not specified). Detection was carried out using a biotinylated secondary antibody (1:200; anti-mouse or anti-rabbit IgG) and Vectastain ABC and DAB kits (Vector Laboratories, Burlingame, CA, USA).

#### Three-Dimensional Organoid Culture

This study was conducted with the approval of the Yonsei University Hospital Institutional Review Board (IRB No. 3-2017-0369), and informed consent was obtained from all tissue donors. Human pancreatic adenocarcinoma specimens were mechanically minced and rinsed three times in DMEM (Welgene, Daegu, Korea) supplemented with 1% FBS and 1× penicillin-streptomycin (P/S). Tissue digestion was performed using 5 mg/mL collagenase II (GIBCO) in DMEM/F12 containing 10 mM HEPES, 1× GlutaMAX, and 1× P/S for 1 hour at 37°C.

Isolated cells were resuspended in organoid culture medium and combined with Matrigel (Corning, MA, USA) at a 1:1 ratio. The mixture was dispensed into 48-well plates and incubated at 37°C for 20 minutes to allow matrix polymerization. Organoids were maintained in a specialized culture medium composed of advanced DMEM/F12 supplemented with 50% Wnt3A-conditioned medium (ATCC, CRL-2647), 10% R-Spondin-1-conditioned medium (Cultrex®), 1× B-27, 10 mM nicotinamide, 10 mM HEPES, 1× GlutaMAX, 1× P/S, 100 ng/mL FGF10 (NKMAX), 1 mM N-acetylcysteine, 50 ng/mL recombinant EGF (PeproTech), 100 ng/mL Noggin (R&D Systems), 500 nM A83-01 (Tocris), 10 nM human [Leu15]-gastrin (Sigma-Aldrich), and 25 µg/mL Plasmocin (InvivoGen). Media were replenished every other day. Organoids were confirmed to be free of mycoplasma contamination.

#### Isolation of Cancer-Associated Fibroblasts (CAFs)

Freshly resected pancreatic tumor tissues were finely chopped and washed in Dulbecco’s phosphate-buffered saline (DPBS) before being plated in 12-well dishes. Cultures were maintained in DMEM with 10% FBS, 1× GlutaMAX, and 1× P/S under standard incubation conditions. CAF identity was confirmed by immunostaining for α-smooth muscle actin and vimentin. Cells between passages 3 and 7 were utilized for downstream assays.

#### Co-Culture of Pancreatic Cancer Organoids and CAFs

Established pancreatic cancer organoids (PCOs) were mechanically disrupted into clusters, while CAFs were dissociated into single cells. For co-culture, 3.3 × 104 PCO cells were mixed with 1 × 105 CAFs (1:3 ratio), resuspended in organoid culture medium, and embedded in Matrigel at a 4:6 ratio. The resulting CAF-integrated PCOs (CIPCOs) were incubated at 37°C with 5% CO2. After 72 hours of drug treatment, cell viability was assessed using the CellTiter-Glo 3D assay (Promega), following the manufacturer’s instructions.

#### Statistical Analysis

Data are presented as mean ± standard error of the mean (SEM). Differences between groups were evaluated using one-way ANOVA followed by Tukey’s multiple comparison test. A p-value of < 0.05 (), < 0.01 (), < 0.001 (), or < 0.0001 (****) was considered statistically significant. All statistical analyses were conducted using GraphPad Prism 9 (GraphPad Software, USA).

## Author Contributions

NSR and SJK equally contributed to this work. GWJ and JHC were involved in conceptualization and writing the manuscript, along with data analysis and interpretation. All authors have read and agreed to the final manuscript.

## Additional information

## Supplementary information

The online version contains supplementary material available

## Correspondence and requests for materials

should be addressed to Jin-Ho Choy or Geun-woo Jin

## Data availability

All other data supporting the findings of this study are availabe in the Supplementary Information.

## Code availability

NA

## References

1 Li, Y. et al. Invasion and metastasis in cancer: molecular insights and therapeutic targets. Signal Transduction and Targeted Therapy 10, 57 (2025). 10.1038/s41392-025-02148-4

2 N, S. R., Jin, G. W. & Choy, J. H. A Strategic Antimetastatic Solution for Bone-Targeting Prostate Cancer via Nanoengineered Niclosamide. Nano Lett (2025). 10.1021/acs.nanolett.5c02826

3 Rejinold, N. S., Jin, G. W. & Choy, J. H. Harnessing Nanohybridized Niclosamide for Precision Mpox Therapeutics. Adv Healthc Mater 14, e2404818 (2025). 10.1002/adhm.202404818

4 Chen, Y. et al. Identification of a novel mechanism for reversal of doxorubicin-induced chemotherapy resistance by TXNIP in triple-negative breast cancer via promoting reactive oxygen-mediated DNA damage. Cell Death & Disease 13, 338 (2022). 10.1038/s41419-022-04783-z

5 Chen, C. et al. Gemcitabine resistance of pancreatic cancer cells is mediated by IGF1R dependent upregulation of CD44 expression and isoform switching. Cell Death & Disease 13, 682 (2022). 10.1038/s41419-022-05103-1

6 Nussinov, R., Yavuz, B. R. & Jang, H. Molecular principles underlying aggressive cancers. Signal Transduction and Targeted Therapy 10, 42 (2025). 10.1038/s41392-025-02129-7

7 Shi, X. et al. Mechanism insights and therapeutic intervention of tumor metastasis: latest developments and perspectives. Signal Transduction and Targeted Therapy 9, 192 (2024). 10.1038/s41392-024-01885-2

8 Wu, B., Zhang, B., Li, B., Wu, H. & Jiang, M. Cold and hot tumors: from molecular mechanisms to targeted therapy. Signal Transduction and Targeted Therapy 9, 274 (2024). 10.1038/s41392-024-01979-x

9 Qin, S. et al. Emerging role of tumor cell plasticity in modifying therapeutic response. Signal Transduction and Targeted Therapy 5, 228 (2020). 10.1038/s41392-020-00313-5

10 Huang, J. et al. Extracellular matrix and its therapeutic potential for cancer treatment. Signal Transduction and Targeted Therapy 6, 153 (2021). 10.1038/s41392-021-00544-0

11 Tomlin, H. & Piccinini, A. M. A complex interplay between the extracellular matrix and the innate immune response to microbial pathogens. Immunology 155, 186–201 (2018). 10.1111/imm.12972

12 Wang, J. et al. Progression from ductal carcinoma in situ to invasive breast cancer: molecular features and clinical significance. Signal Transduction and Targeted Therapy 9, 83 (2024). 10.1038/s41392-024-01779-3

13 Mancini, A. et al. Multiple aspects of matrix stiffness in cancer progression. Front Oncol 14, 1406644 (2024). 10.3389/fonc.2024.1406644

14 Doyle, C. et al. Desmoplasia in cutaneous squamous cell carcinoma with perineural invasion. Clin Exp Dermatol 48, 1279–1280 (2023). 10.1093/ced/llad251

15 Han, X. et al. Reversal of pancreatic desmoplasia by re-educating stellate cells with a tumour microenvironment-activated nanosystem. Nature communications 9, 3390 (2018). 10.1038/s41467-018-05906-x

16 Killock, D. Immunotherapy: Desmoplasia is no barrier to PD-1 blockade in melanoma. Nat Rev Clin Oncol 15, 200–201 (2018). 10.1038/nrclinonc.2018.16

17 Yang, Z. et al. Reprogramming of stromal fibroblasts by SNAI2 contributes to tumor desmoplasia and ovarian cancer progression. Mol Cancer 16, 163 (2017). 10.1186/s12943-017-0732-6

18 Whatcott, C. J. et al. Desmoplasia in Primary Tumors and Metastatic Lesions of Pancreatic Cancer. Clin Cancer Res 21, 3561–3568 (2015). 10.1158/1078-0432.CCR-14-1051

19 Schober, M. et al. Desmoplasia and chemoresistance in pancreatic cancer. Cancers (Basel) 6, 2137–2154 (2014). 10.3390/cancers6042137

20 Lee, J. I. & Campbell, J. S. Role of desmoplasia in cholangiocarcinoma and hepatocellular carcinoma. J Hepatol 61, 432–434 (2014). 10.1016/j.jhep.2014.04.014

21 Merika, E. E., Syrigos, K. N. & Saif, M. W. Desmoplasia in pancreatic cancer. Can we fight it? Gastroenterol Res Pract 2012, 781765 (2012). 10.1155/2012/781765

22 DeClerck, Y. A. Desmoplasia: a response or a niche? Cancer Discov 2, 772–774 (2012). 10.1158/2159-8290.CD-12-0348

23 Dongre, A. & Weinberg, R. A. New insights into the mechanisms of epithelial–mesenchymal transition and implications for cancer. Nature Reviews Molecular Cell Biology 20, 69–84 (2019). 10.1038/s41580-018-0080-4

24 Davoli, E., Zucchetti, M., Giavazzi, R., Garattini, S. & Frapolli, R. Pseudo-resistance to anticancer drugs. Ther Adv Med Oncol 14, 17588359221136776 (2022). 10.1177/17588359221136776

25 Zhao, Y. et al. Stromal cells in the tumor microenvironment: accomplices of tumor progression? Cell Death & Disease 14, 587 (2023). 10.1038/s41419-023-06110-6

26 Rejinold, N. S., Choi, G., Jin, G. W. & Choy, J. H. Transforming Niclosamide through Nanotechnology: A Promising Approach for Long COVID Management. Small 21, e2410345 (2025). 10.1002/smll.202410345

27 Rejinold, N. S., Jin, G. W. & Choy, J. H. Insight into Preventing Global Dengue Spread: Nanoengineered Niclosamide for Viral Infections. Nano Lett 24, 14541–14551 (2024). 10.1021/acs.nanolett.4c02845

28 Rejinold, N. S., Piao, H., Choi, G., Jin, G. W. & Choy, J. H. Niclosamide-Exfoliated Anionic Clay Nanohybrid Repurposed as an Antiviral Drug for Tackling Covid-19; Oral Formulation with Tween 60/Eudragit S100. Clays Clay Miner 69, 533–546 (2021). 10.1007/s42860-021-00153-6

29 Piao, H. et al. Niclosamide encapsulated in mesoporous silica and geopolymer: A potential oral formulation for COVID-19. Microporous Mesoporous Mater 326, 111394 (2021). 10.1016/j.micromeso.2021.111394

30 Rejinold, N. S., Piao, H., Jin, G. W., Choi, G. & Choy, J. H. Injectable niclosamide nanohybrid as an anti-SARS-CoV-2 strategy. Colloids Surf B Biointerfaces 208, 112063 (2021). 10.1016/j.colsurfb.2021.112063

31 Rejinold, N. S., Choi, G., Piao, H. & Choy, J. H. Bovine Serum Albumin-Coated Niclosamide-Zein Nanoparticles as Potential Injectable Medicine against COVID-19. Materials (Basel) 14 (2021). 10.3390/ma14143792

