## supporting files for "Targeted disruption of Oncogenic Biosphere: A Paradigm Shift Beyond Cancer Cell-Centric Therapies"

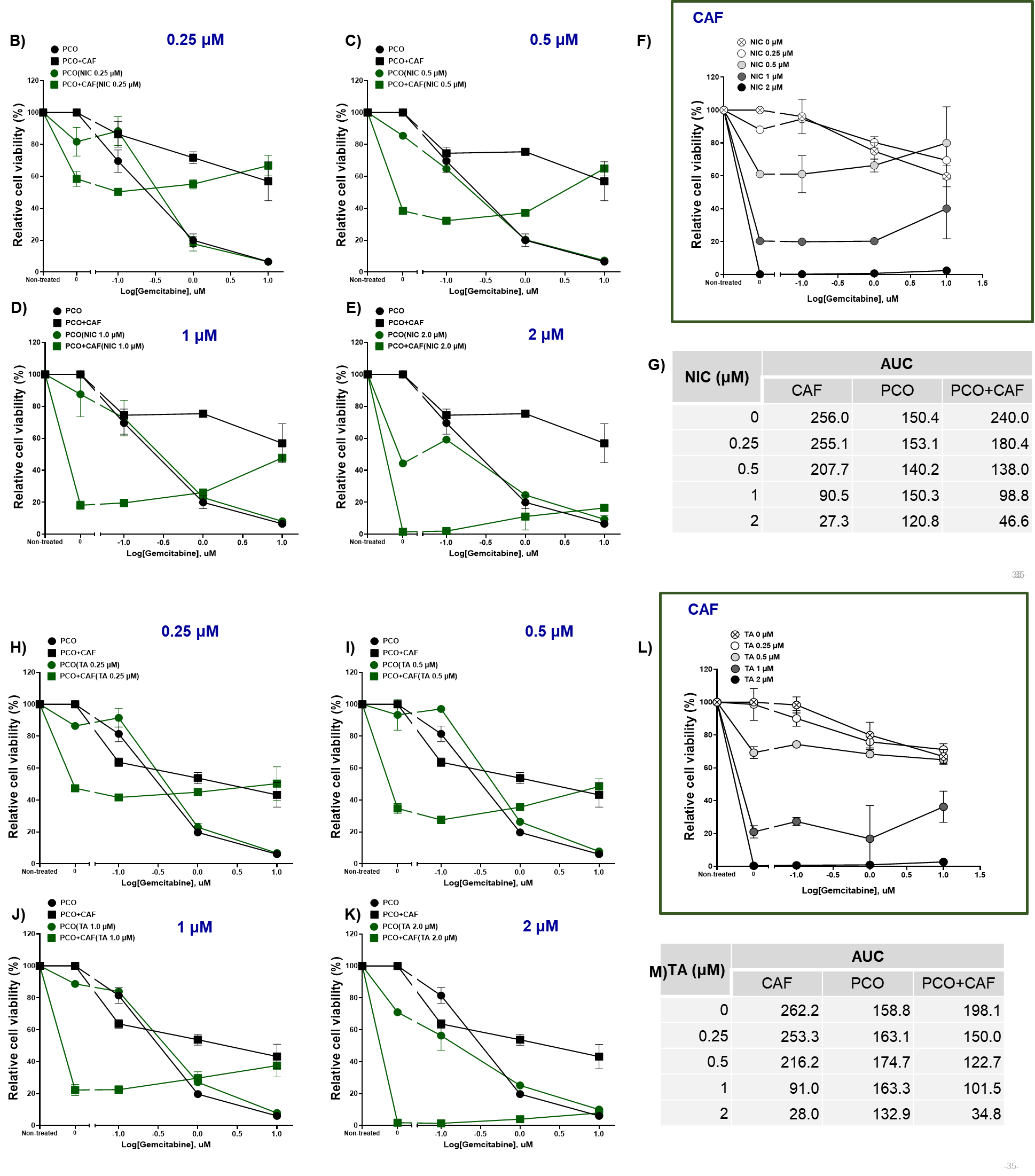

**Figure S 1 | Enhanced dual-compartment cytotoxicity of Penetrium™ versus niclosamide in a second patient-derived pancreatic organoid–CAF co-culture model (hPC 20-001).**

Dose–response analysis of niclosamide (top) and Penetrium™ (bottom) was conducted in patient-derived pancreatic cancer organoids (PCOs), cancer-associated fibroblasts (CAFs), and PCO+CAF co-cultures over 5 days. Viability curves (left) show the effects at 0.25, 0.5, 1, and 2 µM, while AUC-based scatter plots (right) illustrate cumulative cytotoxicity. Corresponding tables present area under the curve (AUC) data for each condition. Both treatments show dose-dependent cytotoxicity, with Penetrium™ demonstrating greater suppression of CAF viability at intermediate concentrations and superior efficacy in the co-culture context. These findings reinforce Penetrium™'s potential for microenvironment-modulating anticancer therapy in desmoplastic tumors.

**Overcoming CAF-Mediated Drug Resistance via ECM-Modulating Nanotherapy**

Despite advances in chemotherapeutics, pancreatic ductal adenocarcinoma (PDAC) remains notoriously resistant to conventional drugs due to its desmoplastic stroma and the presence of cancer-associated fibroblasts (CAFs), which induce extracellular matrix (ECM)-mediated pseudo-resistance. To address this, we performed a side-by-side comparison of free niclosamide and its nanoformulated counterpart, Penetrium™ (NIC–MgO–HPMC), in a clinically relevant organoid model derived from patient tumors (PCO), either alone or in co-culture with CAFs. As shown in **Figure S2**, free niclosamide showed moderate attenuation of gemcitabine efficacy in the presence of CAFs at lower concentrations (e.g., AUC increase from 85.2 to 113.7 at 0.25 μM). While a partial reversal of resistance was noted at higher concentrations (e.g., AUC drop from 58.8 to 20.3 at 2.0 μM), Penetrium™ outperformed across the board. Especially at 2.0 μM, Penetrium™ retained potent cytotoxicity in the CAF-rich environment, reflected by a sharply decreased AUC (from 58.3 in PCO alone to 8.8 in PCO+CAF). This suggests an ECM-modulatory mechanism enabling enhanced drug penetration and possible CAF reprogramming or depletion. These findings corroborate our hypothesis that smart nanoformulations can remodel the tumor microenvironment (TME), reduce the protective barrier conferred by CAFs, and enhance chemotherapeutic sensitivity. Penetrium™, as an ECM-modulating nanodrug, represents a promising translational strategy for tackling dense-stromal tumors like PDAC, where conventional therapies have thus far failed to penetrate the fibrotic barriers effectively.

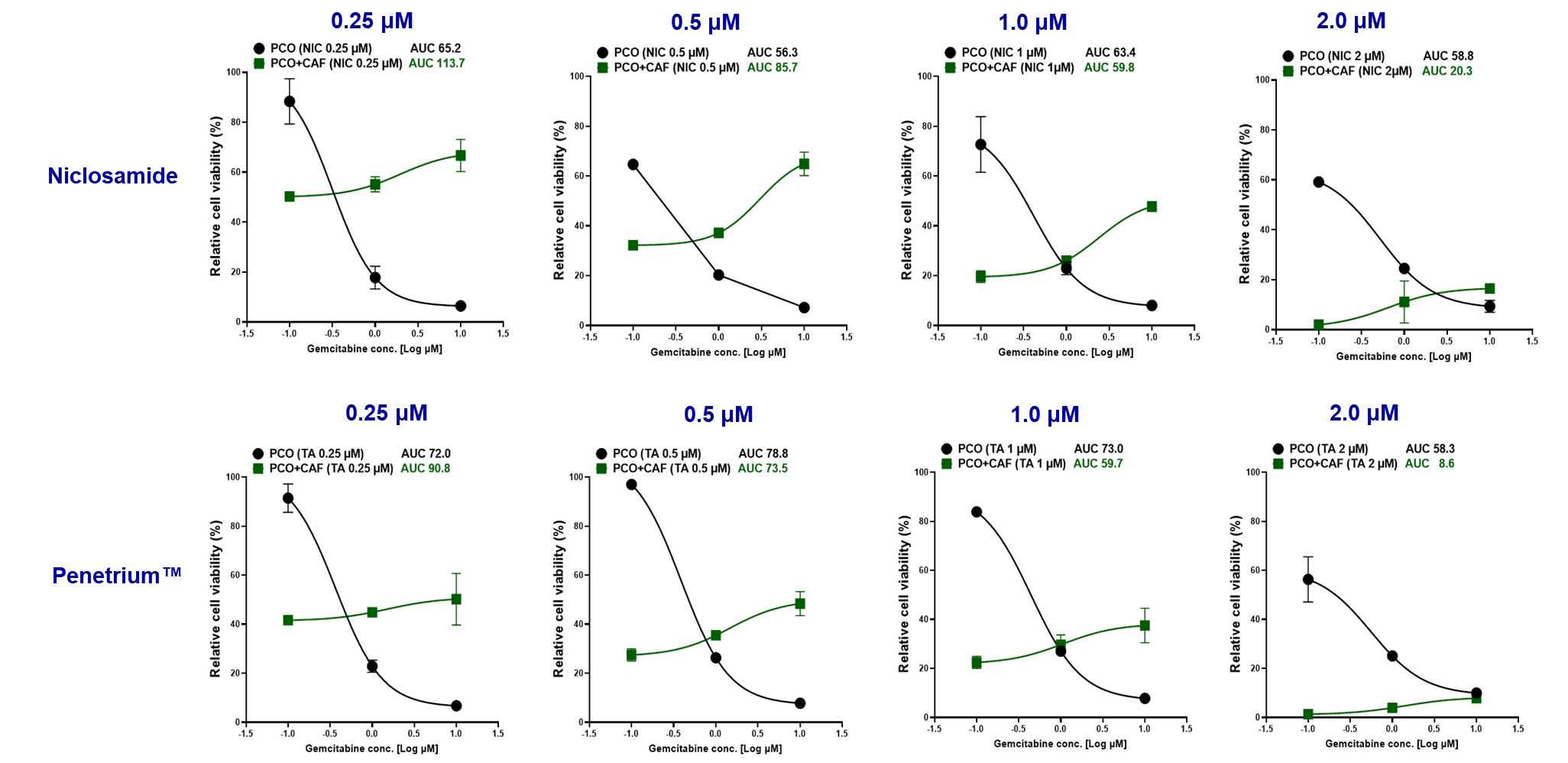

**Figure S 2|Differential anti-tumor efficacy of Niclosamide and Penetrium™ in patient-derived pancreatic cancer organoid (PCO) and CAF co-culture models**

Dose–response curves of gemcitabine sensitivity in the presence of either free niclosamide (top row) or Penetrium™ (niclosamide–MgO–HPMC formulation; bottom row) at four fixed concentrations (0.25, 0.5, 1.0, and 2.0 μM) were evaluated in PCO mono-culture and PCO co-cultured with cancer-associated fibroblasts (CAF). Cell viability was measured after treatment, and area under the curve (AUC) values were calculated to assess the modulatory impact of CAFs on drug efficacy. Notably, Penetrium™ demonstrated a significantly enhanced CAF-suppressive effect at 2.0 μM, with a drastic reduction in AUC (8.8 in PCO+CAF) compared to free niclosamide (AUC 20.3 in PCO+CAF), indicating improved tumor penetration and microenvironmental modulation.

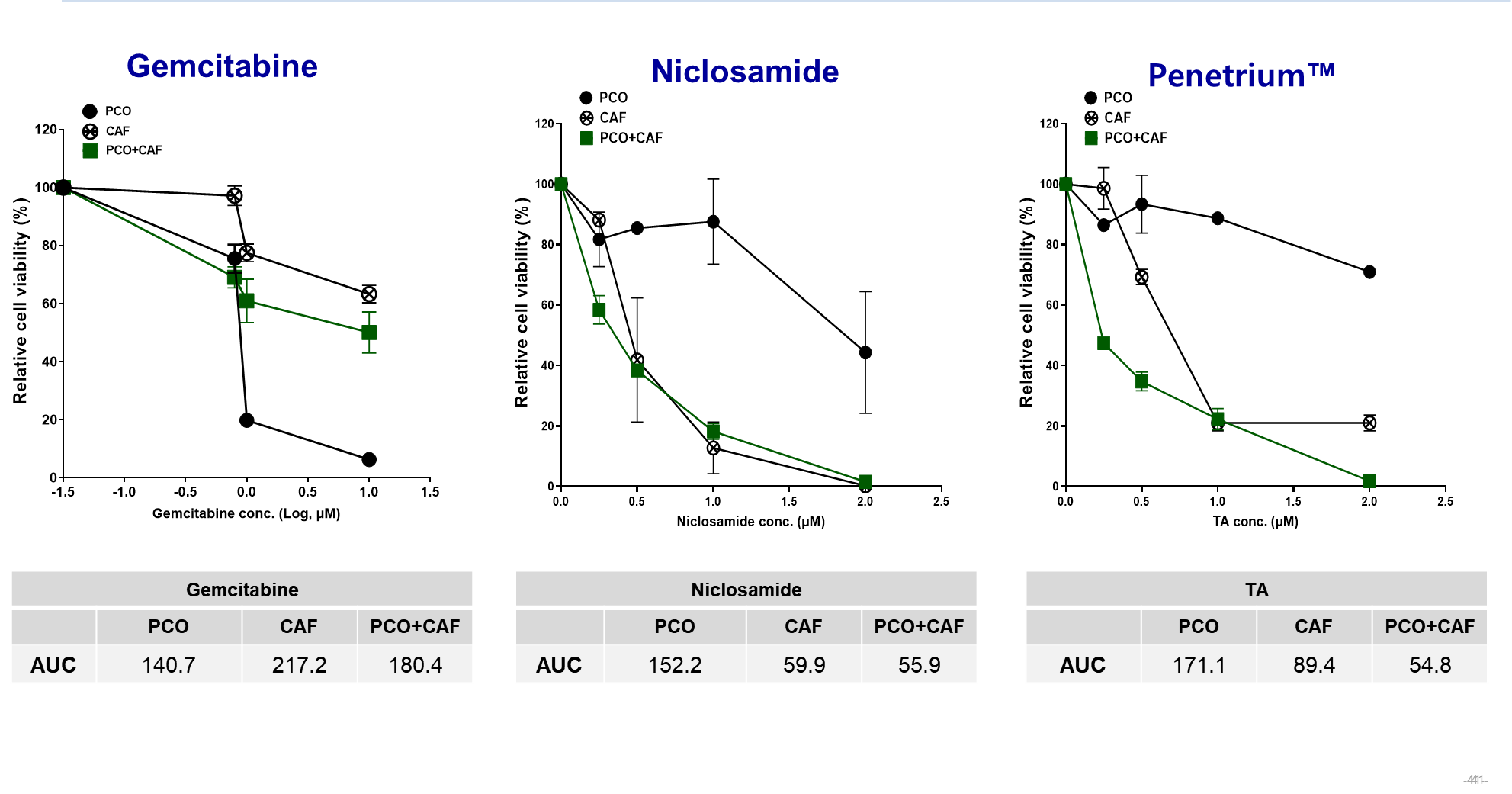

**Figure S 3|** **Comparative cytotoxic effects of gemcitabine, niclosamide, and Penetrium™ in patient-derived pancreatic organoids and CAF co-cultures.**

Dose–response curves of gemcitabine sensitivity in the presence of either free niclosamide (top row) or Penetrium™ (niclosamide–MgO–HPMC formulation; bottom row) at four fixed concentrations (0.25, 0.5, 1.0, and 2.0 μM) were evaluated in PCO mono-culture and PCO co-cultured with cancer-associated fibroblasts (CAF). Cell viability was measured after treatment, and area under the curve (AUC) values were calculated to assess the modulatory impact of CAFs on drug efficacy. Notably, Penetrium™ demonstrated a significantly enhanced CAF-suppressive effect at 2.0 μM, with a drastic reduction in AUC (8.8 in PCO+CAF) compared to free niclosamide (AUC 20.3 in PCO+CAF), indicating improved tumor penetration and microenvironmental modulation.

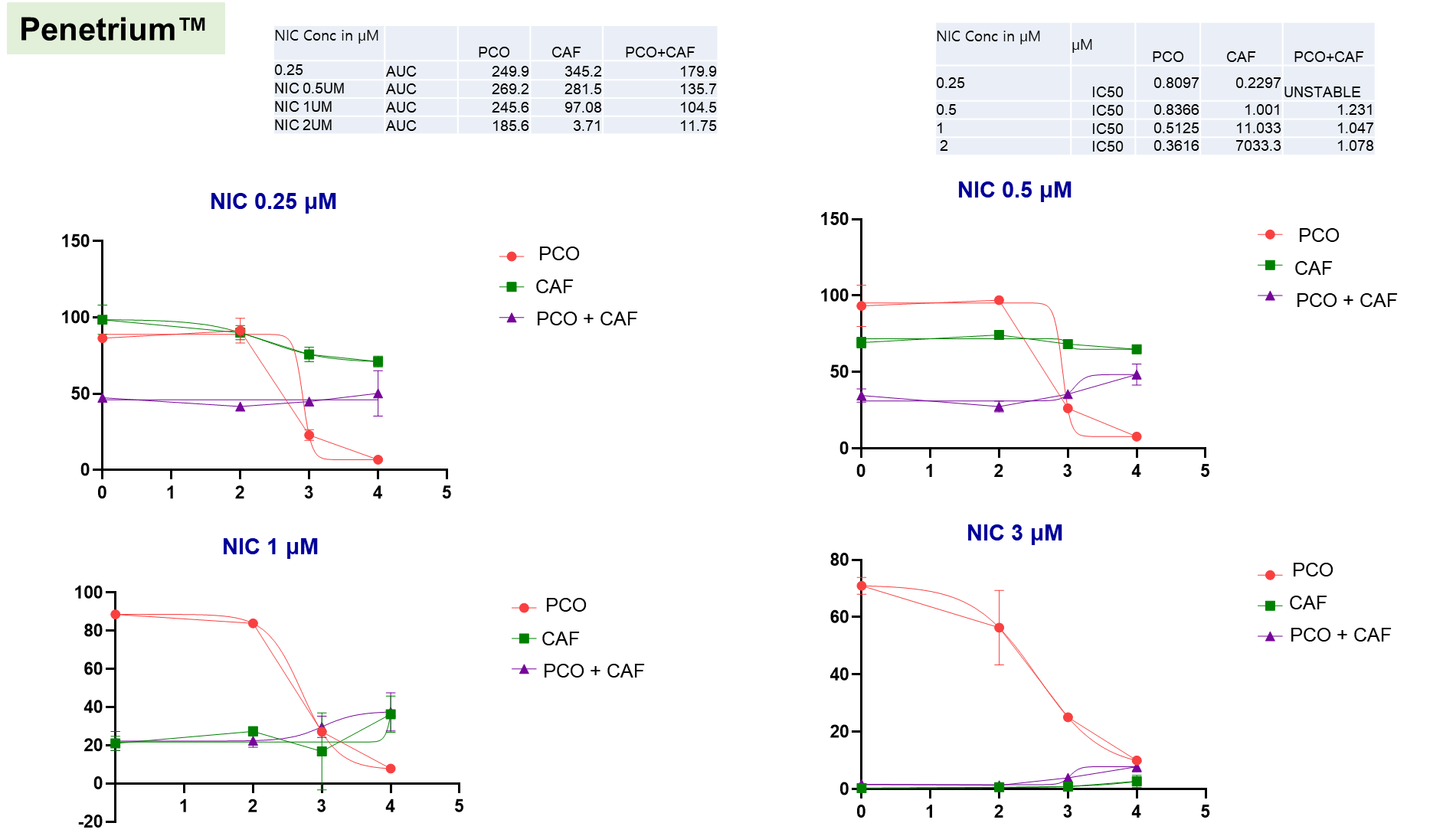

**Figure S 4|** **Stromal-selective cytotoxic modulation by niclosamide at increasing concentrations in pancreatic cancer organoid and CAF models.**

Dose–response viability profiles of gemcitabine were evaluated in the presence of four fixed concentrations of niclosamide (0.25, 0.5, 1, and 3 µM) across three experimental conditions: (i) organoid-only (red), (ii) CAF-only (green), and (iii) organoid+CAF co-cultures (purple). Data shown are plotted on a log scale for gemcitabine concentration. Total peak area (TPA) and IC50 values were calculated and are presented in the accompanying tables. Notably, niclosamide at 2 µM nearly abolished CAF viability (TPA 3.71), while sustaining cytotoxicity in the co-culture (TPA 11.75), suggesting stromal depletion and improved tumor drug accessibility. A similar trend was observed in IC50 shifts, where niclosamide pre-treatment (2 µM) increased gemcitabine IC50 in CAFs to >7000, reflecting stromal collapse. These data collectively support the hypothesis that niclosamide preferentially disrupts CAF survival and mitigates ECM-mediated chemoresistance.

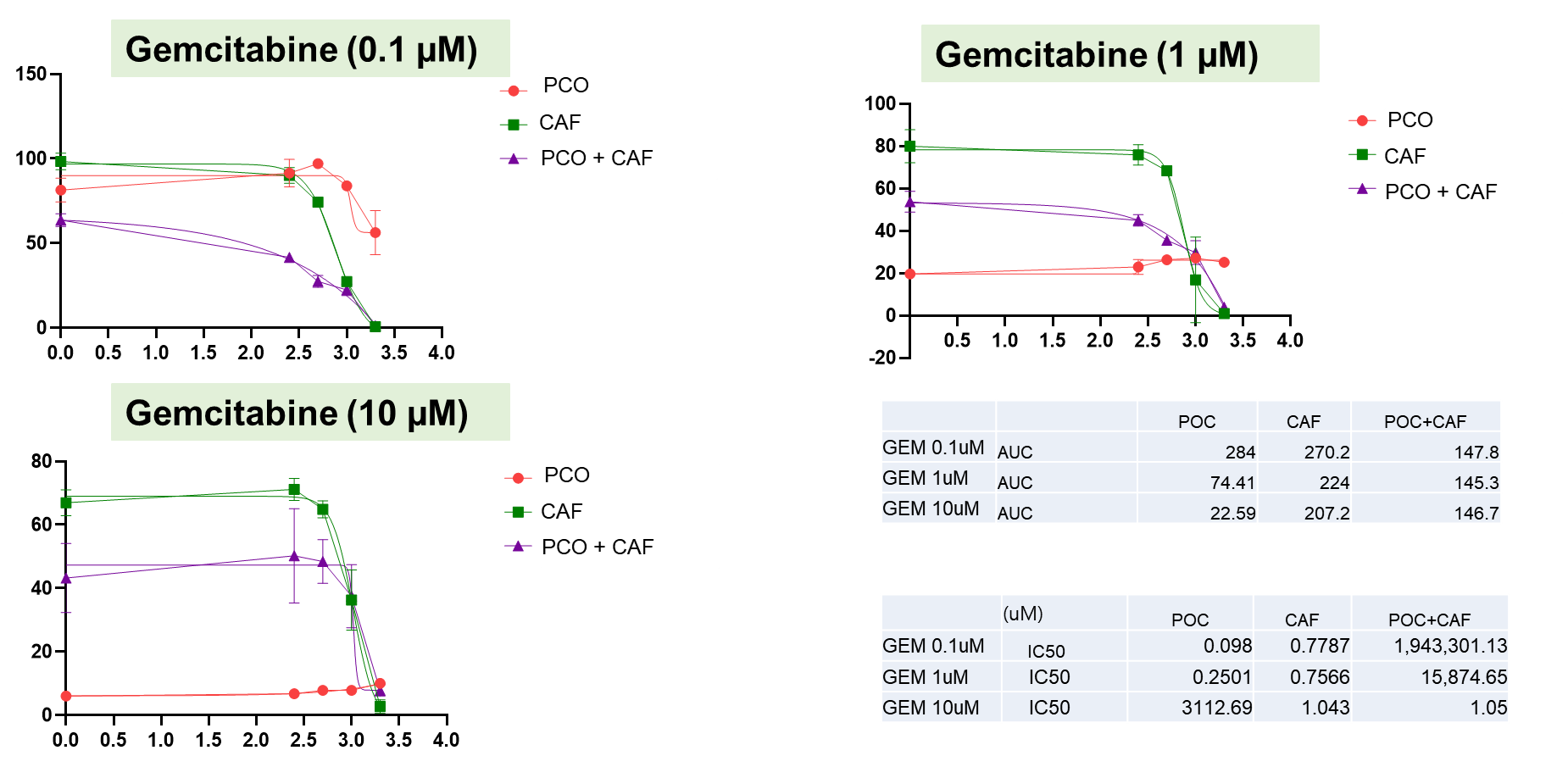

**Figure S 5|** **Stromal barrier effect persists across gemcitabine concentrations in pancreatic tumor organoid models.**

Dose–response viability curves for gemcitabine at 0.1 µM, 1 µM, and 10 µM in three in vitro conditions: pancreatic cancer organoids (PCO; red), CAFs (green), and co-cultured PCO+CAFs (purple). Area under the curve (AUC) and IC50 values are summarized in the accompanying tables. Despite increased gemcitabine dosing, CAF viability remained high (AUC ~207–270), and the co-culture (PCO+CAF) consistently exhibited reduced drug sensitivity, with minimal AUC reduction from 147.8 to 146.7. Notably, IC50 values in the co-culture remained significantly elevated (e.g., 1,943,301.13 at 0.1 µM and 15,874.65 at 1 µM), suggesting stromal shielding. Even at 10 µM gemcitabine, while IC50 in the co-culture dropped (1.05 µM), CAF resistance persisted, reinforcing the need for CAF-suppressive agents to overcome matrix-induced therapeutic evasion.

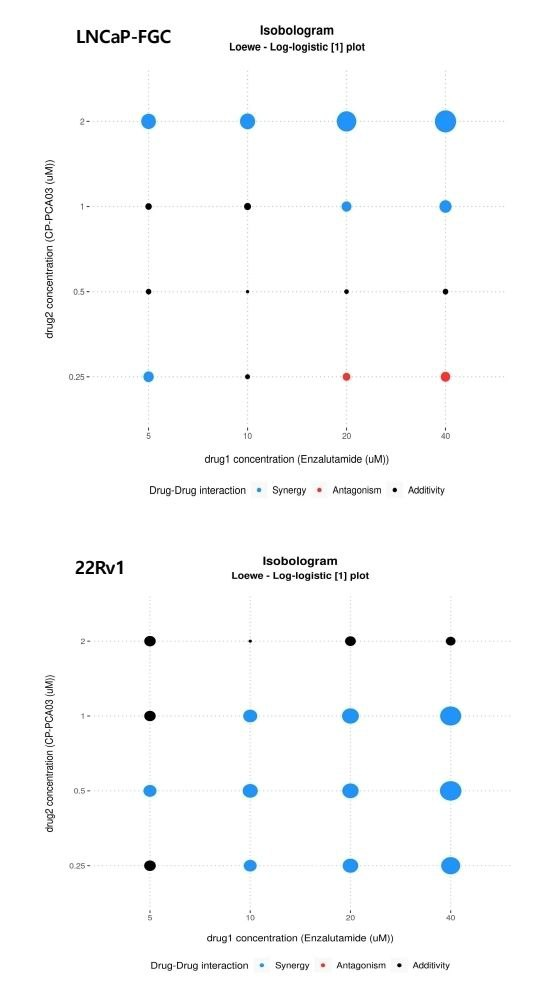

**Figure S 6|** **Drug–drug interaction analysis between Enzalutamide and Penetrium™ in androgen-sensitive and -resistant prostate cancer cell lines.**

Isobolograms were generated using Loewe additivity models to evaluate the combinatorial effects of Enzalutamide (x-axis) and Penetrium™ (y-axis) in LNCaP-FGC (top) and 22Rv1 (bottom) cells. Each dot represents a unique dose pair tested; the interaction outcome is color-coded as: Blue: synergistic interaction, Red: antagonistic interaction,

Black: additive interaction; The diameter of the circles correlates with interaction strength. In LNCaP-FGC cells, synergy was observed predominantly at higher doses of Penetrium™ (1–2 μM) across a broad range of Enzalutamide concentrations. Antagonism was noted at the lowest Penetrium™ concentration (0.25 μM) combined with high Enzalutamide (20–40 μM). In contrast, 22Rv1 cells exhibited consistent synergy across nearly all tested combinations, especially from 0.25–1 μM Penetrium™ with 10–40 μM Enzalutamide, indicating more robust combinatorial efficacy in Enzalutamide-resistant cells.

**Penetrium™** **Synergizes with Enzalutamide in Both Sensitive and Resistant Prostate Cancer Cells**

To evaluate potential synergy between penetrium™ and standard androgen receptor (AR) inhibition therapy, we performed isobologram-based synergy mapping using the Loewe additivity model in LNCaP-FGC (androgen-sensitive) and 22Rv1 (androgen-resistant) cell lines. Dose–response interaction grids were generated across a matrix of Penetrium™(0.25–2 μM) and Enzalutamide (5–40 μM) concentrations (Figure S6). In **LNCaP-FGC**, synergy was observed at higher Penetrium™ concentrations (≥1 μM) across a wide range of Enzalutamide doses. However, lower Penetrium™(0.25 μM) combined with high Enzalutamide (20–40 μM) showed weak antagonism, suggesting dose-dependent interaction variability in sensitive cells. By contrast, **22Rv1 cells** displayed robust synergy across nearly all dose combinations, particularly at mid-range concentrations (0.5–1 μM Penetrium™+ 10–40 μM Enzalutamide). This uniform synergistic effect in the resistant model highlights the potential of Penetrium™as a **sensitizing agent**, potentially overcoming AR-targeted therapy resistance.

These findings support the rationale for **dual therapy approaches**, especially in advanced, Enzalutamide-refractory prostate cancers.
